# Investigating the Consistency of Extracellular Vesicle Production from Breast Cancer Subtypes Using CELLine Adherent Bioreactors

**DOI:** 10.1101/2022.04.11.487804

**Authors:** Colin L. Hisey, Anastasiia Artuyants, George Guo, Vanessa Chang, Gabrielle Reshef, Martin Middleditch, Bincy Jacob, Lawrence W. Chamley, Cherie Blenkiron

## Abstract

Extracellular vesicle (EV) research has grown rapidly in recent years, largely due to the potential use of EVs as liquid biopsy biomarkers or therapeutics. However, in-depth characterisation and validation of EVs produced using conventional *in vitro* cultures can be challenging due to the large area of cell monolayers and volumes of culture media required. To overcome this obstacle, multiple bioreactor designs have been tested for EV production with varying success, but the consistency of EVs produced over time in these systems has not been reported previously. In this study, we demonstrate that several breast cancer cell lines of different subtypes can be cultured simultaneously in space, resource, and time efficient manner using CELLine AD 1000 systems, allowing the consistent production of vast amounts of EVs for downstream experimentation. We report an improved workflow used for inoculating, maintaining, and monitoring the bioreactors, their EV production, and the characterisation of the EVs produced. Lastly, our proteomic analyses of the EVs produced throughout the lifetime of the bioreactors show that core EV-associated proteins are relatively consistent, with few minor variations over time, and that tracking the production of EVs may be a convenient method for indirectly monitoring the bioreactors’ health. These findings will aid future studies requiring the simultaneous production of large amounts of EVs from several cell lines of different subtypes of a disease and other EV biomanufacturing applications.

## Introduction

Extracellular vesicles (EVs) are subcellular membrane-bound particles that contain several types of biomolecules indicative of their parental cells. Once released from the parental cell, they can be found in all bodily fluids and can be internalised by both nearby and distant cells to influence their functions. The recent exponential growth in EV research has validated that this shuttling of EVs and their cargo between cells plays a key role in pathological and physiological processes [1-3]. In addition, their inherent biocompatibility makes them promising candidates as therapeutic delivery vehicles following isolation from desirable donor cells, such as mesenchymal stem cells [4-6] or following modifications of their surface and contents [7-9]. Perhaps the most promising use of EVs is as biomarkers in liquid biopsy applications, particularly in diseases such as cancer [10-12]. However, the low concentration of cancer-associated EVs in patient samples and difficulty producing them in abundance using cell lines grown in physiologically relevant conditions has somewhat hindered progress in this area.

To circumvent this obstacle, several commercial and custom-built bioreactor systems have been established as viable alternatives to using extensive numbers of conventional tissue culture flasks [13]. Many of these systems have been repurposed from hybridoma antibody production applications, as the molecular weight cut-offs used to retain antibodies are also applicable to large and small EVs. They have all demonstrated the ability to significantly improve EV yields by up to ∼100 fold by increasing cell density, as well as providing several more-physiologically relevant features such as growth surface properties and dynamic fluid interactions [14, 15].

Besides simply increasing total EV production, they provide several other advantages such as significantly reducing the total amount of single use plastic used, providing highly concentrated EVs, which reduces or eliminates the need for downstream concentration steps, and typically do not require passaging of cells or other extensive time and reagent commitments. Several disadvantages include the inability to dynamically image the cultured cells due to the opacity or shape of the growth surfaces, the moderate cost of setup and media consumption, the risk of contamination and failure, an incomplete understanding of their effect on cell behaviour relative to conventional cultures, and in some cases, the need for additional equipment such as pumps.

One promising bioreactor design used for EV production is hollow-fiber bioreactors, for example the Fibercell system, where one of several types of fibre bundle designs (5-20 kDa) are contained within a cylindrical cartridge and media is continually pumped through the porous fibres while adherent or suspension cells are cultured in the volume outside of the fibres [3]. While this bioreactor requires a pump system, it has been reported to produce significantly greater numbers of EVs per volume compared to conventional culture flasks. It has been primarily used for anti-cancer or regenerative medicine therapeutic applications by producing EVs from mesenchymal stromal cells [16-20] or engineered HEK293 cells [21-23]. Another interesting and related design is a 3D-printed perfusable bioreactor that was used to produce EVs from endothelial cells and to determine whether ethanol conditioning could improve the bioactivity of the isolated EVs [24]. Clearly, perfusable bioreactors are useful for producing large quantities of EVs, but the requirement of pump systems and significant incubator space may not be ideal if several cell lines need to be cultured simultaneously.

A simpler and potentially more accessible bioreactor design is the CELLine AD 1000 (adherent) static two chamber bioreactor, where a small, high-density cell chamber is separated from a larger media chamber by a 10 kDa semi-permeable, cellulose acetate membrane. In this system, no pump is needed as gas exchange occurs passively through a bottom silicone membrane while nutrient and waste exchange occur passively through an upper 10 kDa membrane. EVs produced by the cultured cells are concentrated in the 15 mL of conditioned media in the cell chamber. These CELLine AD 1000 (adherent) and CL 1000 (suspension) flasks have been used to culture several types of cancer cell lines for a wide range of downstream applications and were first shown to be effective for improving EV production from both mesothelioma and natural killer (NK) cells [25]. Improvements have since been made in the culture media formulations by utilising advanced media to minimise bovine EV contamination while maintaining high levels of cell viability [26]. Several other studies have used similar culture conditions and focused on the downstream RNA and proteomics analyses which are enabled by the production of abundant EVs using these reactors [27-33]. One recent study demonstrated the culture of non-malignant HEK-293 cells and subsequent EV isolation, loading with siRNA, and uptake by pancreatic cancer cells [34]. Importantly, another recent study using prostate cancer cell lines and metabolomic analyses demonstrated that the cells are grown in a more starved environment compared to flat plastic, with limited access to selected nutrients [35]. This, however, is more physiologically relevant for cancer microenvironments and therefore should mimic their *in vivo* growth more closely.

In breast cancer, EVs have been implicated in playing important roles in disease progression and metastasis, immune evasion, inhibiting treatment efficacy, and have demonstrated significant potential as liquid biopsy biomarkers [36-40]. However, due to the aforementioned issues in producing large amounts of breast cancer EVs *in vitro*, progress has been hindered for using cell line-derived EVs to ultimately improve patient outcomes. In this study, we report the successful long-term growth of five cell lines that represent the breast cancer disease spectrum using CELLine AD 1000 bioreactor flasks. We present a reliable approach for inoculation, maintenance, and monitoring of the cultures, followed by efficient EV isolation. We also show the growth structure of each cell line within the bioreactors presenting as diverse 3D, tissue-like structures based on SEM and histological imaging. We also validate EV collections based on nanoparticle tracking analysis (NTA), transmission electron microscopy (TEM), and western blotting, and demonstrate minimal variation in core EV-associated proteins using mass spectrometry proteomics at different time points. As this field continues its rapid growth, these findings will provide confidence to other researchers who desire to continually produce an abundance of consistent EVs from several cell lines simultaneously for biomarker, therapeutic, or other EV-related applications.

## Materials and Methods

### Bioreactor Inoculation, Adaptation, and Maintenance

*Media A*: DMEM (Gibco) with 10% fetal bovine serum (FBS, Merck) and 1% Penicillin/Streptomycin (PS, Gibco). *Media B*: Advanced DMEM/F-12 (Gibco), 2% CDM-HD (Fibercell), 2% FBS, 1% Glutamax (Gibco), and 1% PS. *Media C*: Advanced DMEM/F-12, 2% CDM-HD, 1% Glutamax (Gibco), and 1% PS.

Several breast cancer cell lines across the spectrum of the disease, classified by the presence of estrogen receptor (ER), progesterone receptor (PR), and human epidermal growth factor receptor 2 (HER2) [27], including MCF7 (ER+/PR+/HER2-), BT-474 (ER+/PR+/HER2+), SKBR3 (ER-/PR-/HER2+), BT-20 (ER-/PR-/HER2-), and MDA-MB-231 (ER-/PR-/HER2-) [41] were seeded at a minimum of 1.5 × 10^6^ cells/mL in 15 mL Media A into the lower cell chamber of a CELLine AD 1000 bioreactor flask (Argos) and with 500 mL of the same media in the upper media chamber (**Fig. 1A**). Cells were gradually adapted to CDM-HD serum replacement (Fibercell Systems) and Advanced DMEM/F12 (to minimise FBS usage) by substituting Media C for Media A. The cell chamber was refreshed every 3-4 days, and the media chamber was refreshed every week. After one week, media was changed to 75% Media A and 25% Media B. The following week it was changed to 50% Media A and 50% Media B, then the following week to 25% Media A and 75% Media B. Finally, in week 4, the cell chamber was carefully washed 3x with 15 mL prewarmed PBS to remove any residual FBS EVs and filled with 15 mL Media C, while the media chamber was maintained with 500 mL Media B. This approach allowed the continual use of FBS in the media chamber without bovine EVs passing through the 10 kDa membrane and contaminating the cell-derived EVs (**Fig. 1B**).

**Figure 1.**
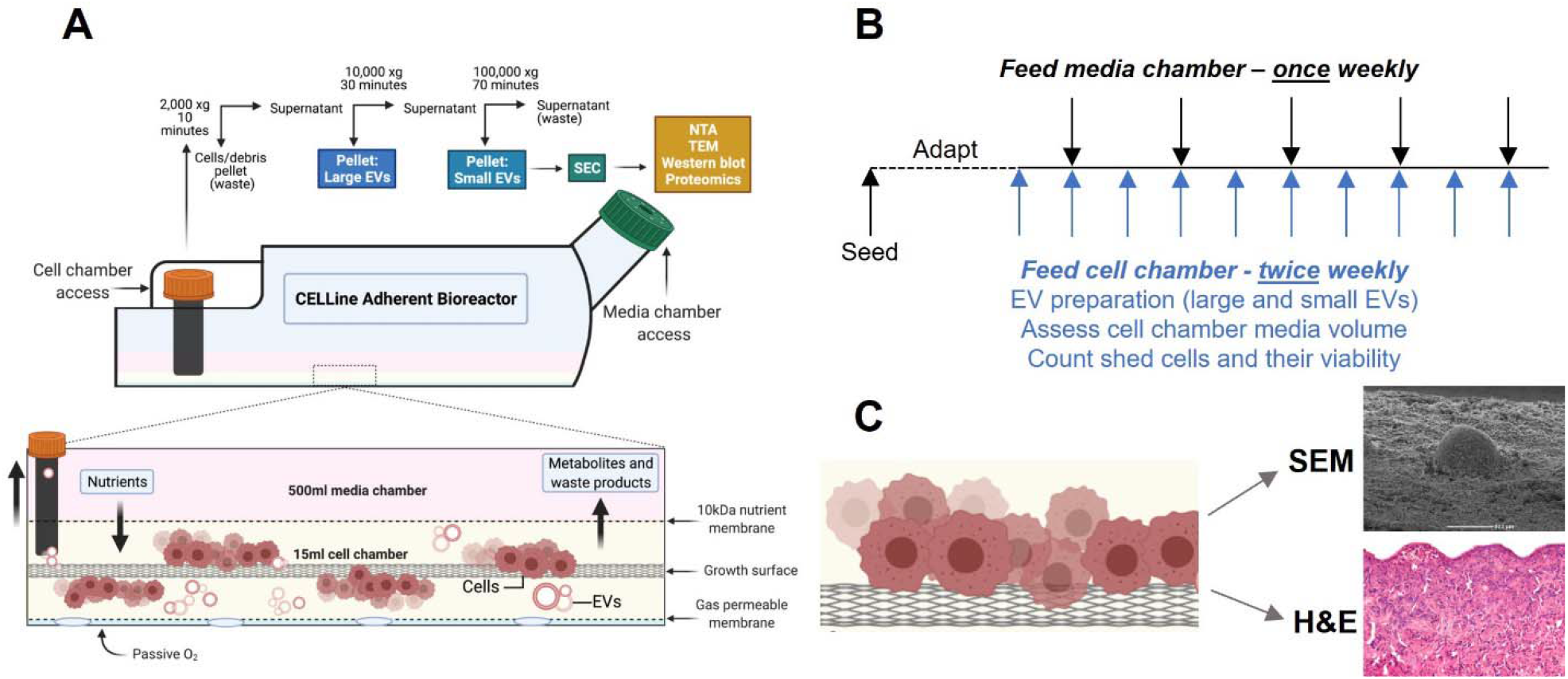
Schematic of bioreactor experimentation and maintenance. (A) Diagram of two-chambered CELLine AD 1000 bioreactor and process flow for isolating and characterising EVs; (B) Bioreactor maintenance approach showing EV collections twice per week following media adaptation, and media chamber refreshment once per week; (C) Methods of imaging bioreactor growth surface following deconstruction including SEM and H&E. Images created in biorender.com.

Following adaptation, 15 mL of conditioned media from the cell chamber was harvested for EV isolation and refreshed every 3-4 days (Monday and Thursday), while the media chamber was changed every 7 days (Thursday). The cell chamber media volume (mL) was noted at each replacement. 10 µL of the complete conditioned media was used for quantitation of suspension cells by combining it with 1:1 Trypan blue and counting on a Countess automated cell counter (Invitrogen). Bioreactors were maintained until greater than 20 mL of media was removed from the cell chamber on consecutive harvests, indicating that the dialysis membrane was damaged (BT-474-1 and MDA-MB-231). A further indicator of a failed bioreactor was abrupt, heavy cell loss in the shed material (MCF7). Two bioreactors were halted in March 2020 due to COVID-19 interruption (BT-20 and SKBR3). The final two bioreactors, replicates of the BT-474 cell-line (BT-474-2 and BT-474-3), were halted once sufficient EVs were collected for planned experiments. In all cases, the reactors were washed thrice with PBS and fixed in 4% glutaraldehyde in PBS at 4°C overnight.

### Conventional Cultures for Comparison

BT-474 and MDA-MB-231 cells were cultured serum-starved in 2D flasks to compare the EV yields to bioreactor cultures. Briefly, cells were cultured in DMEM with 10% FBS and 1% P/S and were bulked up to 9 x T175 flasks of each by splitting and reseeding during their growth phase (∼80% confluence) using TrypLE (Gibco). The 9 x T175 flasks were seeded at ∼30% confluence in serum-containing media, and once the cells reached ∼60% confluence, they were washed 3x with PBS to remove any residual exogenous FBS EVs, and 50 mL of DMEM with 1% P/S was added to each flask. After 48h, this conditioned media was collected and centrifuged at 2,000 xg for 10 min, followed by concentration using a Vivaspin 50, 100 kDa concentrator to reduce the media volume from 450 mL down to 40 mL. EVs from this concentrated conditioned media were ultracentrifuged and purified using size exclusion chromatography (SEC; see below for methods).

Both conventional cell cultures and cultures grown in bioreactors, were tested for mycoplasma using Mycoplasma Stain Kit (Sigma-Aldrich) prior to seeding.

### EV Isolation

The 15 mL of conditioned media from the cell chamber of the bioreactor was centrifuged at 2,000 xg for 10 min to remove cells and other debris. The supernatant was then centrifuged at 10,000 xg for 30 min (JA.30-50 Ti rotor, Avanti, Beckman Coulter) to pellet the large EVs (also known as microvesicles) and this pellet was resuspended in 500 µL PBS and stored. The supernatant was then ultracentrifuged at 100,000 xg for 70 min (JA.30-50 Ti rotor, Avanti, Beckman Coulter) to yield a crude small EV pellet. This pellet was resuspended in 700 µL PBS and stored at -80^0^C until needed, minimising the number of freeze thaws.

500 µL of crude small EVs were loaded onto a 35 nm qEV Original SEC column (Izon Science Ltd.), and fractions 7 through 23 were collected using an automated fraction collector (Izon Science Ltd, 500 µL per fraction). A high sensitivity BCA assay (Pierce, ThermoFisher Scientific) was performed for each collected fraction to determine their protein concentration. Once EV-rich fractions were determined, only those EV-rich fractions (F8-F10) were collected and pooled in subsequent isolations.

### Nanoparticle Tracking Analysis

SEC-purified small EV individual and pooled fractions were diluted in PBS at a 1:100 ratio and measured with a NS300 Nanosight (Malvern Analytical). Three 30 sec videos were taken under low flow conditions (Screen gain: 2, Camera level: 9) and characterised using the Nanosight 3.0 software (Screen gain: 4, Detection threshold: 4) to calculate mean and mode particle diameters, concentration, and size distribution. EV-rich fractions were pooled and used for tracking EV production, TEM, and mass spectrometry. Statistical comparisons of average EV production from each cell line were performed in Graphpad Prism using an ordinary one-way ANOVA with Tukey’s multi-comparison test to determine statistically significant differences between groups (p<.05). Averages are reported with a ± standard error mean.

### Transmission Electron Microscopy

Negative staining TEM of small EVs was conducted by adsorption onto Formvar-coated copper grids (Electron Microscopy Sciences) for 10 min. Excess liquid was removed with filter paper (Whatman), and then the copper grid was transferred to 20 µL of filtered 2% uranyl acetate for 2 min. Excess liquid was removed with filter paper, and the grid was allowed to dry for 10 min. Grids were visualised on Tecnai G2 Spirit TWIN (FEI, Hillsboro, OR, USA) TEM at 120 kV accelerating voltage. Images were captured using a Morada digital camera (SIS GmbH, Munster, Germany).

### Scanning Electron Microscopy

Glutaraldehyde-fixed CELLine AD 1000 flasks were deconstructed using a Dremel tool from the bottom side by cutting around the growth surface area to access and cut up the dialysis membrane (upper), growth surface (middle), and gas permeable membrane (bottom). Sections of these layers were washed with PBS for 2 × 5 min, followed by a series of dehydration treatments of 35% ethanol for 10 min, 50% ethanol for 2 × 10 min, 70% ethanol for 2 × 10 min, 90% ethanol for 2 × 10 min, and 100% ethanol for 4 × 10 min. Samples were sputter coated with 10 nm of gold at 1nm/min (Q150R S, Quorum) and imaged using a JCM-6000 benchtop SEM (Jeol) at 15 kV.

### Hematoxylin and Eosin Staining

Glutaraldehyde fixed cells on the growth surface were dehydrated and embedded in paraffin wax end on and in flat orientation. 5 µm sections were floated onto histobond slides, air dried and then stained for hemotoxylin and eosin (H&E) after standard de-paraffinisation. Due to the use of glutaraldehyde for fixation, immunostaining of these sections was not possible.

### Immunoblotting

Pooled small EVs were concentrated by ultracentrifugation at 100,000 xg for 70 min, and protein levels were measured by BCA. Equal protein quantities (4µg/lane) of SEC-purified small EVs and cell lysates were separated by SDS-polyacrylamide gel electrophoresis (PAGE; NuPAGE 4-12% Bis-Tris Protein Gel, ThermoFisher Scientific) and transferred to PVDF membranes (Millipore) using a Trans-Blot Turbo Transfer System (BioRad). To prevent nonspecific binding, membranes were incubated with blocking solution - 5% non-fat milk powder in PBS-T (0.05% vol/vol Tween20 in PBS) for 1 hour. Membranes were immunoblotted with CD9 (ab92726, 1:1000), CD63 (sc-5275, 1:20,000), CD81 (ab79559, 1:1000), GRP94 (ab3674, 1:1000), GAPDH (ab9485, 1:5000), EpCAM (ab71916, 1:1000), ERα (ab108398, 1:1000) antibodies, followed by either biotinylated anti-mouse (115-065-071, 1:10,000) or rabbit (111-065-046, 1:10,000) secondary IgG antibodies (JacksonImmuno Research), and finally HRP-conjugated streptavidin (016-030-084, JacksonImmuno Research, 1:10,000). Bound antibody was visualised using Pierce™ ECL Western Blotting Substrate (ThermoFisher Scientific), and the chemiluminescence was measured using BioRad ChemiDoc MP imaging system.

### Label-Free Proteomic Analysis (SWATH-MS)

EVs were concentrated to a volume of 150 µL in a vacuum concentrator (ThermoSavant, Holbrook, NY, USA). After the addition of 150 µL of 7 M urea, 2 M Thiourea, 5mM DTT, 0.1% SurfactAmps X-100 in 50 mM ammonium bicarbonate, the samples were sonicated in a sonic bath for 15 min. Disulphide bonds were reduced by incubation at 56°C in a heat block for 20 min. Cysteines were alkylated by the addition of iodoacetamide (IAM) to 15 mM final concentration and incubated in the dark at room temperature for 30 min, followed by the addition of cysteine to 15 mM final concentration in order to quench residual IAM.

For the SWATH analysis, 15 µg of total protein was taken from 15 higher concentration samples and 5 µg for the other 3 samples (BT-474, later fractions) which had very low total protein concentrations. Each sample was then diluted 10-fold with 50 mM ammonium bicarbonate for digestion with 0.375 µg (for 15 µg protein samples) or 0.125 µg (for 5 µg protein samples) sequencing grade modified porcine trypsin (Promega, Madison, WI, USA). The samples were incubated at 45°C for 1 hour in a chilled microwave (CEM, Matthews, NC, USA) using 15W power. An additional 0.375 µg of trypsin (0.125 µg for the three low protein samples) was then added to each sample, followed by incubation overnight at 37°C. After digestion, samples were acidified to pH 3 via addition of 50% formic acid, centrifuged for 3 min at 16,000 xg, and desalted using 10 mg Oasis HLB SPE cartridges (Waters, Milford, MA, USA) as per the manufacturer’s instructions. Purified peptides were eluted with 300 µL of 50% acetonitrile in 0.1% formic acid and then concentrated to a final volume of 25 µL in a vacuum concentrator.

For Ion Library generation, a pooled digest (3 µg of protein from the fifteen 15 µg samples) was applied to an SCX MicroSpin column (The Nest Group, Inc.) according to manufacturer’s instructions and fractionated using 50 mM, 70 mM, 85 mM, 100 mM, 150 mM, 175 mM, 200 mM, 300 mM, 400 mM and 1000 mM NaCl. The collected fractions were desalted on 10 mg Oasis HLB SPE cartridges and vacuum concentrated to 20 µL. LC-MS/MS analysis of these fractions was performed on an Eksigent 425 nanoLC chromatography system (Sciex, Framingham MA, USA) connected to a TripleTOF 6600 mass spectrometer (Sciex). Samples were injected onto a 0.3x 10mm trap column packed with Reprosil C18 media (Dr Maisch) and desalted for 5 min at 10 µL/min before being separated on a 0.075 × 200 mm picofrit column (New Objective) packed in-house with Reprosil 3u C18-AQ media. The following gradient was applied at 300 nL/min using a NanoLC 400 UPLC system (Eksigent): 0 min 5% B; 45 min 35%B ; 47 min 95% B; 50 min 95% B; 50.5 min 5% B; 60 min 5% B where A was 0.1% formic acid in water and B was 0.1% formic acid in acetonitrile.

The picofrit spray was directed into a TripleTOF 6600 Quadrupole-Time-of-Flight mass spectrometer (Sciex) scanning from 300-2000 m/z for 150 ms, followed by 30 ms MS/MS scans on the 60 most abundant multiply-charged peptides (m/z 80-1600) for a total cycle time of ∼2 sec. The mass spectrometer and HPLC system were under the control of the Analyst TF 1.7 software package (Sciex).

SWATH analysis was conducted using an acquisition method comprising 75 variable width isolation windows (with 1 Da overlap) covering a precursor mass range of 400–1100 m/z. The accumulation time was 150 ms for the initial TOF-MS scan and 20 ms for each SWATH MS/MS scan, giving a total cycle time of ∼1.7 s, using the same LC conditions as described above. The resulting peptide fragment ion peak areas were calculated using Skyline software.

The resulting data were searched against a database comprising the human proteome (downloaded from Uniprot, 03 Nov 2020) appended with a set of common contaminant sequences (75,224 entries in total) using ProteinPilot version 5.0 (Sciex). Search parameters were as follows: Sample Type, Identification; Search Effort, Rapid; Special factors, Urea Denaturation, Cys Alkylation, Yes; Digestion, Trypsin; FDR analysis, Yes. The resulting group file exported from ProteinPilot was transferred to Skyline for use as an Ion Library for SWATH analysis.

All data were analysed using the Skyline software and MSStats [41]. The spectral library built from the previous step was attached to the project. A list of targeted protein fasta sequences was imported via the transition rules and spectral library information to the project to generate the target transitions. A retention time calculator was created using a set of the representative peptides. The SWATH results were then read to the project according to the target transitions and SWATH isolation scheme. The peak elution profiles were extracted within the +/- 3 minutes window of the predicted retention times. The protein data is summarized via Tukey’s median polish method on all transitions and normalized via equalizing medians based on reference signals to remove the inter-sample error. Missing values were replaced by imputation according to an accelerated failure model. The processed protein features were log2 transformed for heatmap plotting and group comparison analysis. For hierarchical clustering, normalised intensities were first z-scored and then clustered using Euclidean as a distance measure for column and row clustering.

Vesiclepedia (Version 4.1, 2018) and Exocarta (Version 5, 2015) were downloaded to map previously reported EV proteins. Gene ontology (GO) analyses were performed with Database for Annotation, Visualization and Integrated Discovery (DAVID, Version 6.8).

## Results

### Bioreactor Monitoring

Due to the structure of the CELLine AD 1000 bioreactor, it is not possible to view the health or status of the cells besides inspecting for media colour (phenol red pH indicator, which is unreliable according to the manufacturer[42]), bubbles in the cell chamber, or analysis of the contents of the conditioned media once removed from the chamber. While glucose levels have been used in past studies as a means of monitoring [20, 43, 44], media volume, shed cells and EVs can be more easily monitored throughout the life of the bioreactor. Due to the natural turnover of cells on the growth surface and a limited live cell capacity [42], we hypothesised that shed cells could act as a surrogate marker of the bioreactor’s health. Shed cells were counted and assessed for viability during each cell chamber media collection to determine if there were any apparent patterns related to bioreactor health or correlations to EV production (**Fig. 2**). Most cell lines decreased the total number of shed cells over time, with only MCF7 cells exhibiting consistent shedding throughout the life of the bioreactor until a sudden increase in cell number was observed at day 158, resulting in its termination. Shed cell viability is dynamic and unpredictable, with MDA-MB-231 cells slowly decreasing over time, BT-20 cells slowly increasing over time, and the other cell lines with no apparent trend, particularly obvious in the three replicate BT-474 bioreactors.

**Figure 2.**
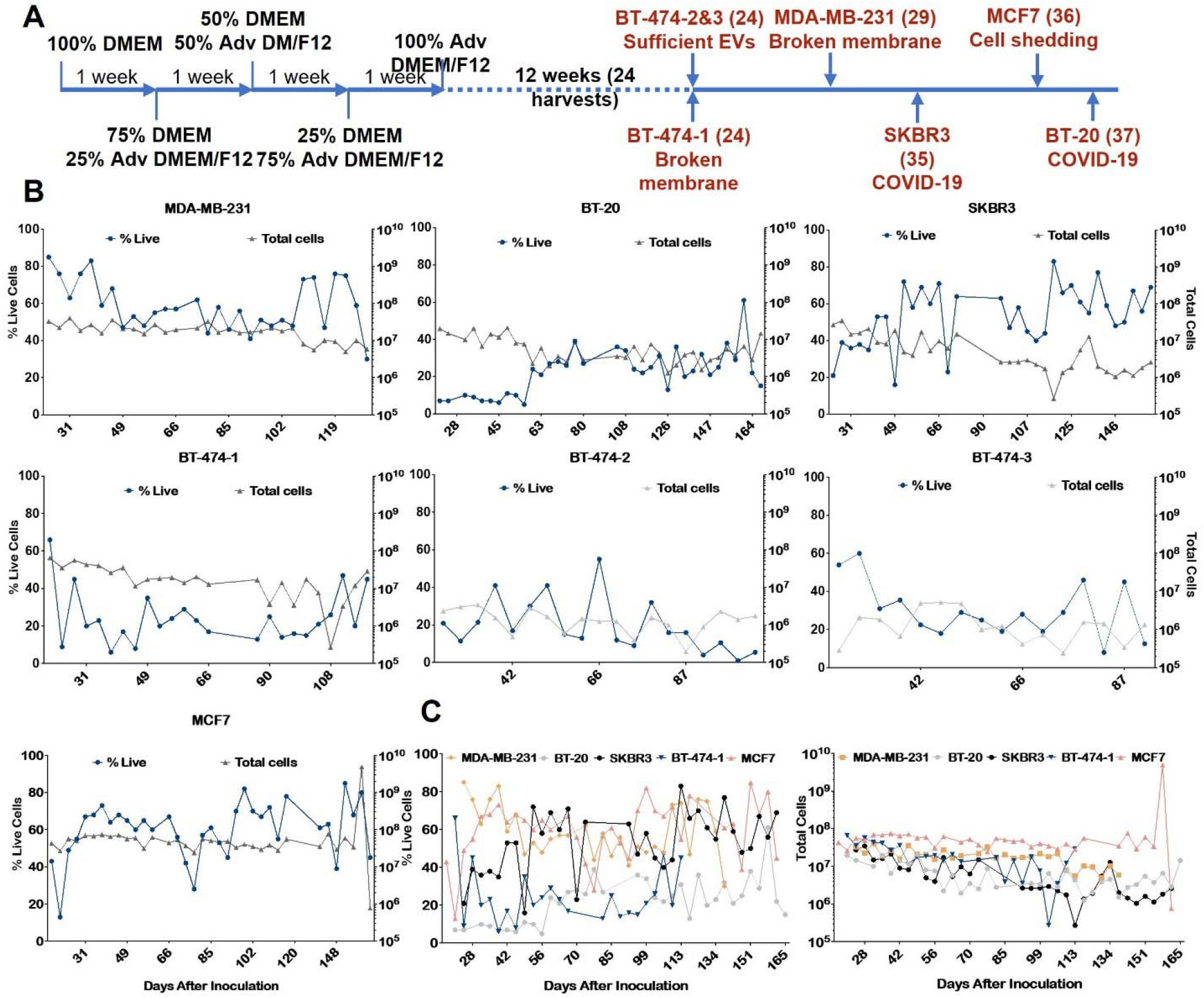
(A) Timeline of CELLine AD 1000 bioreactor experiments. The blue line shows a bioreactor lifetime, with the first four weeks of adaptation followed by at least twelve weeks of sample harvest. Shown in red are the cell lines grown in bioreactors, in parenthesis the number of EV preparations collected and reasons for bioreactors termination; (B) Tracking shed cells (black, right y-axis) and their viability (blue, left y-axis) for individual cell lines throughout the bioreactor lifetimes (days after inoculation). The percentage of viable cells was calculated by Trypan Blue exclusion assay. Each dot represents a single harvest; (C) Viability (left) and total number of shed cells (right) for all five cell lines throughout the bioreactor lifetimes for comparison.

### Extracellular Vesicle Characterisation

Following isolation using SEC, EVs were characterised according to the MISEV guidelines [45]. Small EV size distributions based on NTA measurements were consistent, as expected given the inherent size selection from SEC (Suppl. Fig. 1). In addition, negative staining of the isolated small EVs showed expected EV morphologies (**Fig. 3A**). Western blot showed that EV lysates were enriched for tetraspanins CD81, CD9 and CD63 when compared with whole cell lysates (**Fig. 3B**) although interesting differences in abundance for these three proteins were seen between cell lines. Immunostaining for endoplasmic reticulum protein GRP94 demonstrated purity of the EV preparations. Assessment of two breast cancer-associated markers, demonstrated that small EVs from all cell lines were highly enriched for surface receptor EpCAM (Epithelial Cell Adhesion Molecule) when compared with cell lysates. EpCAM was, as expected, less abundant in the triple-negative, mesenchymal line MDA-MB-231. The nuclear receptor ERα (Estrogen Receptor alpha) was enriched in EVs isolated from MCF7 and BT-474 cell lines similar to the cell lysates, representative of their hormone receptor positive status. Together these analyses highlight the validity of the EV preparations and that they reflect their cell of origin for molecular subtype.

**Figure 3.**
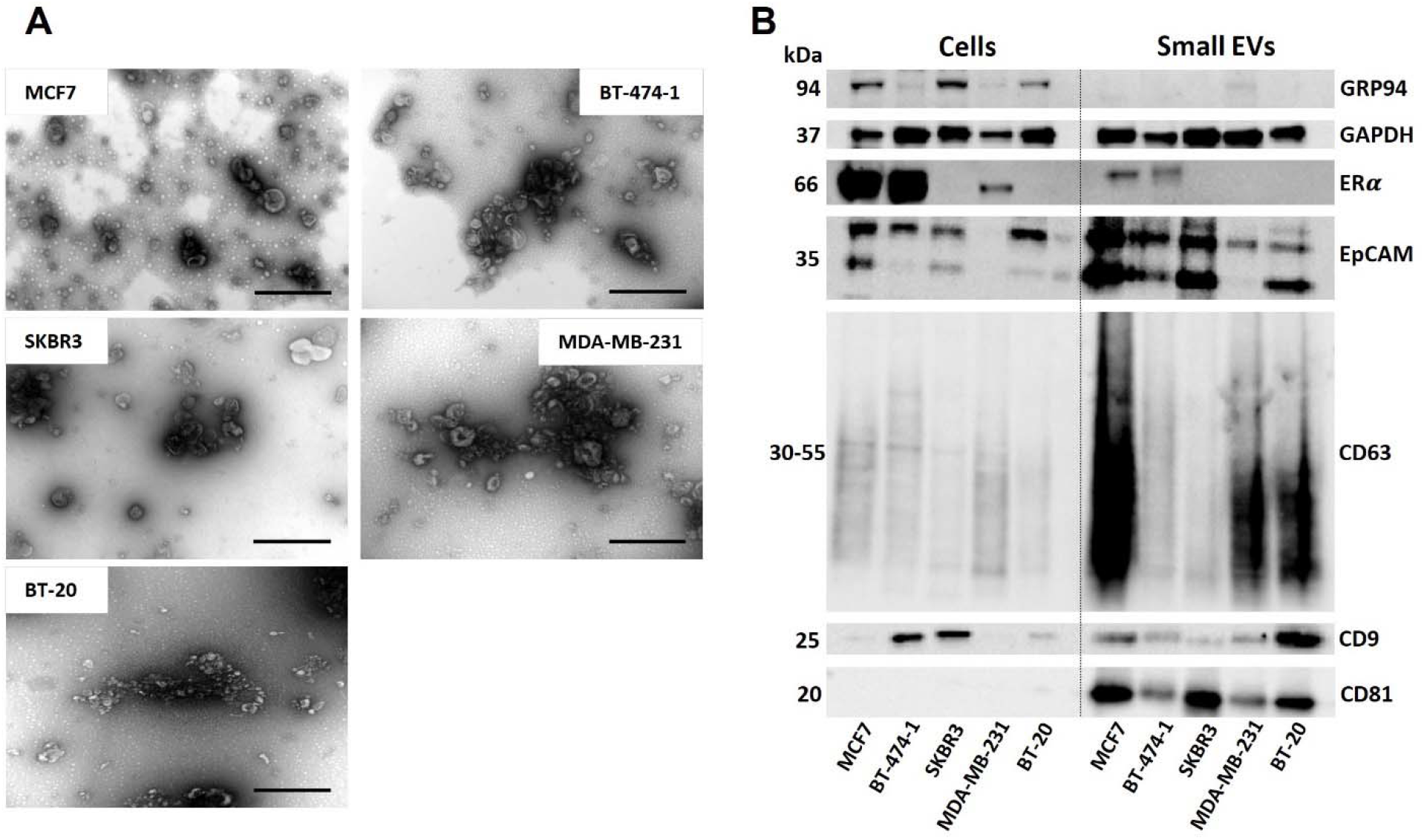
(A) Representative TEM images of small EVs isolated from all bioreactors. Scale bar 400 nm; (B) Western blot results showing enrichment of CD81, CD9 and CD63 in SEC-purified small EVs, as a pool of 6 preparations taken from the lifetime of each bioreactor, compared to the enrichment of GRP94 in cell lysates. Probing with two breast cancer-associated markers - ERα and EpCAM was also performed for small EVs and whole cell lysates. Each lane was loaded with an equal amount of protein (4 µg) based on BCA assay and loading confirmed by probing with an anti-GAPDH antibody. Full blots can be found in Suppl. Fig. 2.

Once EV-rich SEC fractions were identified (F8-F10) and pooled, the particle counts and protein yields over bioreactor lifetimes were plotted. These results demonstrated that while some cell lines slowly increase EV particle counts and EV-protein over time (MDA-MB-231, BT-20, and MCF7), the two HER2+ cell lines (SKBR3, BT-474) showed strikingly sharp decreases in yield after the first few weeks of harvest (**Fig. 4A**). These patterns in EV production were not reflected in the shed cell number or viability (**Fig. 2**).

**Figure 4.**
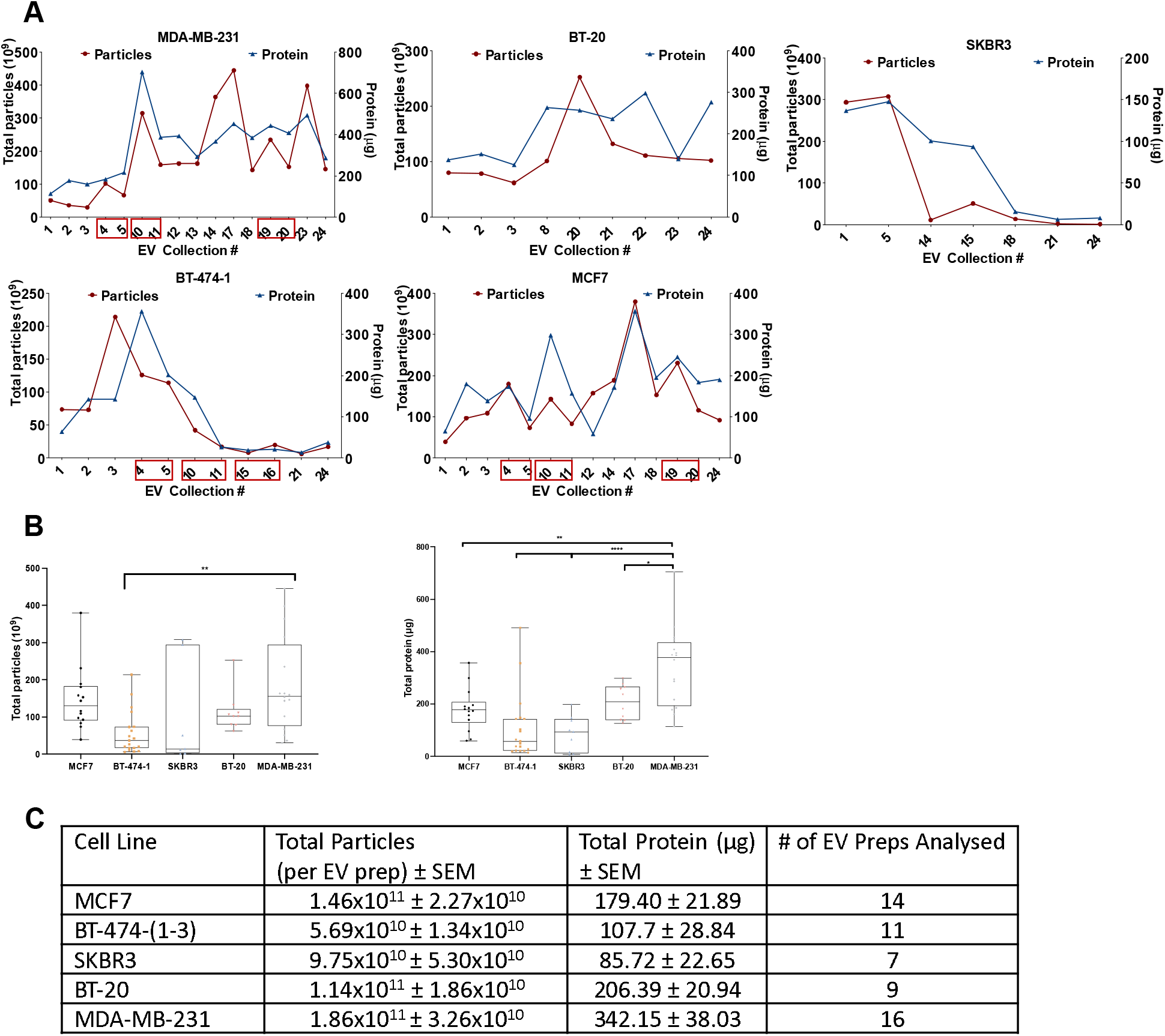
(A) Production of EVs and protein yields for each individual breast cancer cell line in CELLine AD 1000 bioreactors. The total particles number (x10^9^) measured by NTA is shown in red (left y-axis). The protein amount in µg, as assessed by BCA assay, is shown in blue (right y-axis). Each dot represents individual EV prep. Numbers in red boxes indicate EV preps taken for proteomic analysis; (B) Production of EVs for all cell lines in CELLine AD 1000 bioreactors demonstrates a significant difference between MDA-MB-231 and BT-474-(1-3) bioreactors (** p<0.01) and production of EV-protein for all cell lines in CELLine AD 1000 bioreactors demonstrates a significant difference between MDA-MB-231 and BT-20 bioreactors (* p<0.03), MDA-MB-231 and BT-474-1 and SKB3 bioreactors (*** p<0.0001), MDA-MB-231 and MCF7 bioreactors (** p<0.001); (C) Average particles and EV-protein production with standard error mean (SEM) from all bioreactors tested.

Comparison between the average particle counts from each cell line indicated that the only significant difference was between MDA-MB-231 and BT-474-(1-3) bioreactors (p<0.01 **Fig. 4B**), while significant differences were found in EV-protein quantities between MDA-MB-231 and all other cell lines (p<0.0001 **Fig. 4B**). While the calculated average particle counts were quite different, no other statistically significant differences were found, most likely due to the high variance in EV production over the lifetime of several of the bioreactors. The average particle count and EV-protein per preparation were generous for each cell-line, highlighting the value of the bioreactors for production of EV material (**Fig. 4C**).

### Comparison to Conventional Cultures

Comparing the SEC fractions for particle number and protein concentrations between bioreactor and conventional flask-produced small EVs (9x T175 flasks) in serum-free conditions demonstrated that bioreactor-produced EVs were not only much more abundant per harvest but also contained more protein per EV than those from conventional cultures (**Fig. 5)**. For MDA-MB-231 cells, the average harvest of 18.6 × 10^10^ ± 3.26 × 10^10^ EVs from bioreactors compared to 1.27 × 10^10^ EVs from 9x T175 flasks represents a 14.65x average increase in EV production or the rough equivalent of 132x T175 flasks per single bioreactor harvest. For BT-474 cells, the average harvest of 5.69 × 10^10^ ± 1.34 × 10^10^ EVs from bioreactors compared to 4.86x 10^9^ from 9x T175 flasks represents an 11.36x average increase in EV production or roughly 102x T175 flasks per bioreactor harvest. While these increases in yields seem staggering, the incredible increase in cell density as demonstrated by later imaging results, combined with the inherent EV loss that can occur during concentration of the large volume of media from conventional flasks, both likely contribute to this substantial difference.

**Figure 5.**
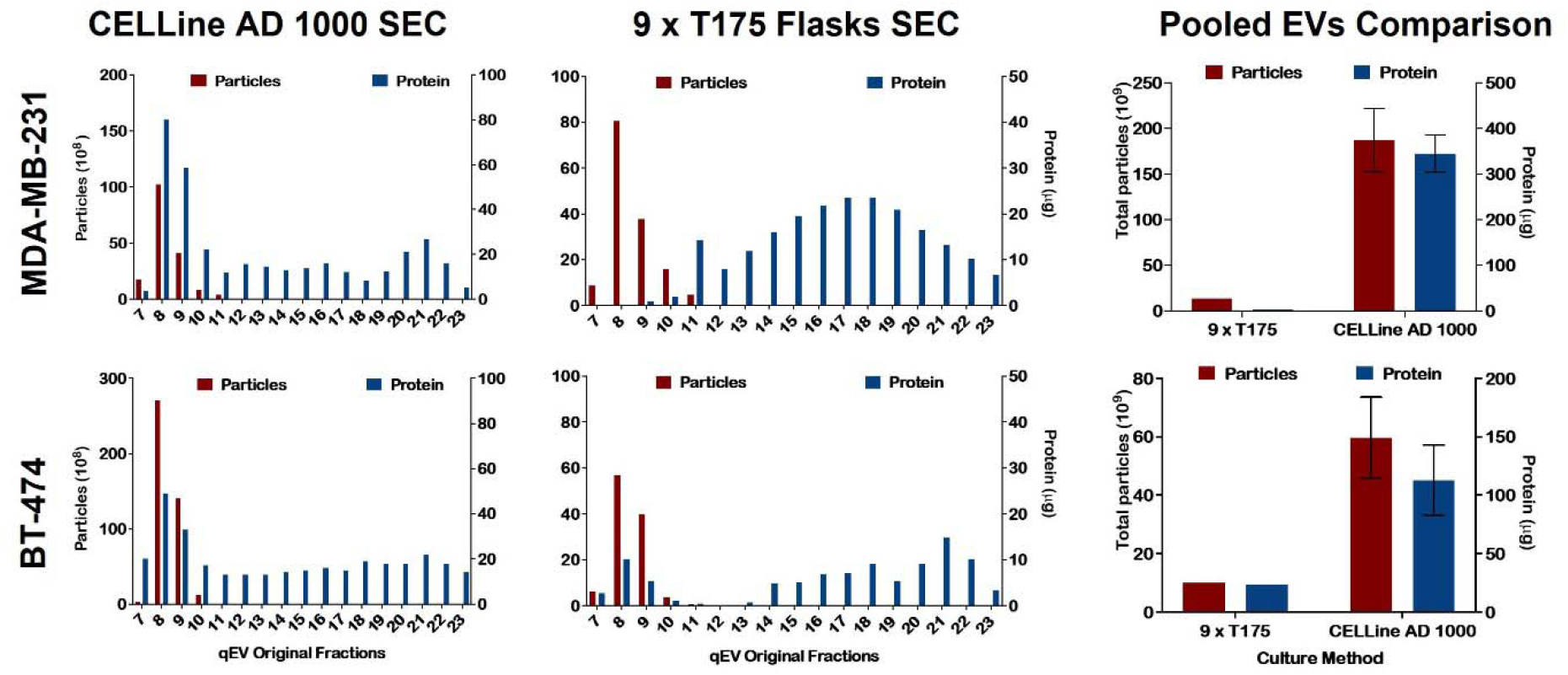
Comparison of CELLine AD 1000 produced EVs and EVs produced in conventional tissue cultures for MDA-MB-231 and BT-474 cell lines. SEC fractions graphed for particle number (x10^9^) based on NTA (red, left y-axis) and protein (µg) by BCA assay (blue, right y-axis). Pooled EVs (SEC fractions 8-10) are compared for total particle and protein yield for multiple bioreactor preparations (error bars are SD) compared to a single 9x T175 flask EV preparation.

### Bioreactor Growth Surface Imaging

To further elucidate the underlying factors which induce EV production within the bioreactors, the growth surfaces were imaged using scanning electron microscopy. On the bare growth surface (no cells), a tightly woven polyethylene terephthalate (PET) fibrous mesh is seen, which is less visible in the cell seeded bioreactors (**Fig. 6**). These types of microscale fibrous structures are known to improve EV production counts from MDA-MB-231 cells [14], but to nowhere near the degree seen in this study. For many of the bioreactors, there were clear, patchy areas of higher cell density that could be seen with the naked eye. Interestingly, the cell masses were thicker for the triple-negative cell lines (BT-20, MDA-MB-231), and the underlying woven mesh is more visible for the three hormone receptor positive cell lines where the cell mass is thinner. Varying degrees of extracellular matrix were also evident. While only qualitative, these density differences fit with the overall trend of small EV production, which was higher for BT20 and MDA-MB-231 cell-lines (**Fig. 4C**). In one case, for the BT-20 bioreactor, spheroid-like structures (∼500 µm) were present on the top side of the growth surface (Suppl. Fig. 3), which was not seen in any other imaged surfaces or cell-line bioreactors.

**Figure 6.**
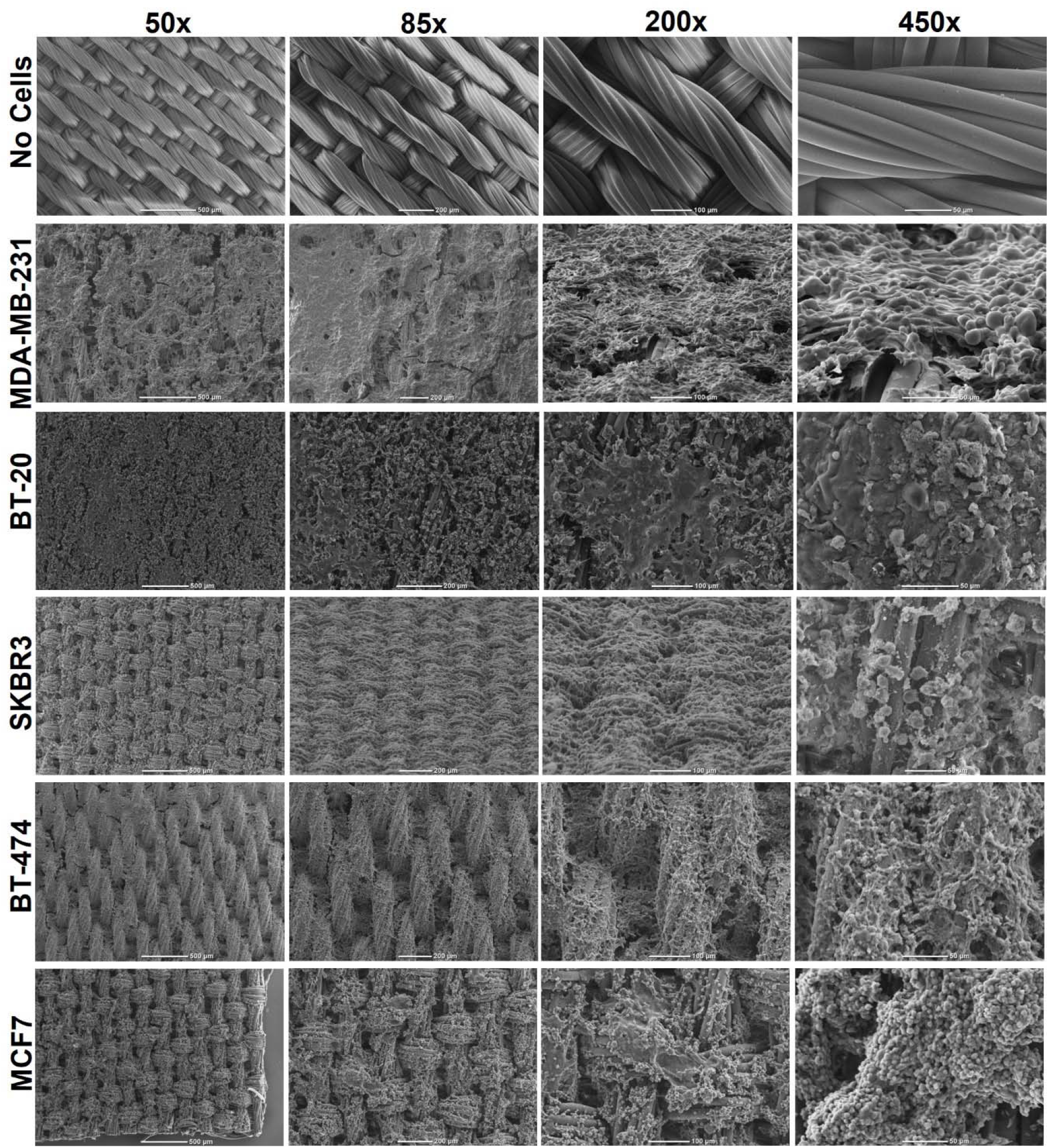
Representative SEM images of all bioreactor growth surfaces showing the bare growth surface, as well as different cell thicknesses and morphologies. Scale bars 500, 200, 100 and 50 µm from left to right in vertical columns. Thick cell masses can be seen for both triple-negative cell lines, (MDA-MB-231 and BT-20) with thinner and more patchy cells for the three hormone receptor-positive cell lines below.

While SEM imaging provided valuable information on the surface structure of the cells on the growth surfaces, it could not provide detailed images of the cell structures within the cell mass. To explore this, H&E sections from BT-20 and BT-474 bioreactors (chosen by the feasibility of timing with bioreactor discontinuation and COVID-19 laboratory shutdowns) were generated. These demonstrated the thick cell mass structures within the bioreactors and their orientation relative to the fibrous growth surfaces (Suppl. Fig. 4), with the BT-20 cells forming organised structures within the main cell mass. Cell growth around the bioreactor surface fibres was also obvious.

### Longitudinal Proteomics of Bioreactor EVs

Several studies have used CELLine AD 1000-produced EVs for proteomic analyses, however none have reported the consistency of EV-protein cargo over the course of the bioreactor lifetimes. This is a key question to ensure reproducibility of the EV preparations. To explore this, six EV harvests from three cell lines (MCF7, BT-474, and MDA-MB-231) were chosen at different time points during bioreactor culture to assess any changes to EV cargo during the lifetime of the bioreactor (Suppl. Data). These included two early (4-6 weeks post-inoculation), two mid (7-9 weeks), and two late (12-14 weeks) for each cell line. In addition, they were chosen as three pairs from harvests within the same week to determine if there were differences between samples collected on media chamber feeding days (Thursday - collected a week after media replacement) compared to 4 days after media chamber replacement (Monday). We hypothesised that there might be EV protein cargo alterations due to cell stress caused by the week-long build-up of waste products and depletion of nutrients in the ‘Thursday’ EVs.

SWATH label-free quantitative proteomics analysis of the 18 EV preparations identified 2,875 proteins present in at least one sample. The top 100 most abundant proteins within each cell line were cross-referenced against the 100 most commonly reported EV associated proteins in Vesiclepedia and Exocarta [46, 47] (Suppl. Fig. 5 and Suppl. Data). Half of the top 100 proteins in each online database were also identified in one or more of the three breast cancer cell lines’ EVs, with a third of the top 100 in either Vesiclepedia or Exocarta present in all datasets, including Alix and HSP90. We also identified several proteins unique to specific cell lines (e.g. Her2/ERBB2 in BT-474, Galtectin-3 in MCF7, and CD44 and Vimentin in MDA-MB-231) that were not commonly reported in the EV databases. Despite the overlap with the databases, tetraspanins CD9, CD81, and CD63 were lacking in our bioreactor EV proteomics data although they were, however, present in trial non-quantitative shotgun proteomics (data not shown) and Western blots (**Fig. 3B**) highlighting known challenges in generating fully representative proteomics data using this methodology [48].

Next, we applied a protein intensity cut off (>20,000) in order to identify proteins that were highly abundant in all 18 of our samples. From this, 1,191 out of the 2,875 unique proteins were labelled as “core” proteins. Gene ontology (GO) analysis using the DAVID database [49] (**Fig. 7**) identified that these proteins were significantly enriched for proteins associated with selected cellular components including ‘extracellular exosome’, ‘membrane’, ‘cytosol’ and ‘cell-cell adherens junction’. The enriched biological process terms for these core EV proteins included many cancer-relevant physiological and pathological processes, including ‘cell-cell adhesion’, ‘mRNA splicing’, and ‘intracellular protein transport’. Finally, the GO terms for molecular function of the core EV proteins were enriched for ‘protein binding’, ‘cadherin binding involved in cell-cell adhesion’, and ‘poly(A) RNA binding’ (full list in Suppl. Data Table).

**Figure 7.**
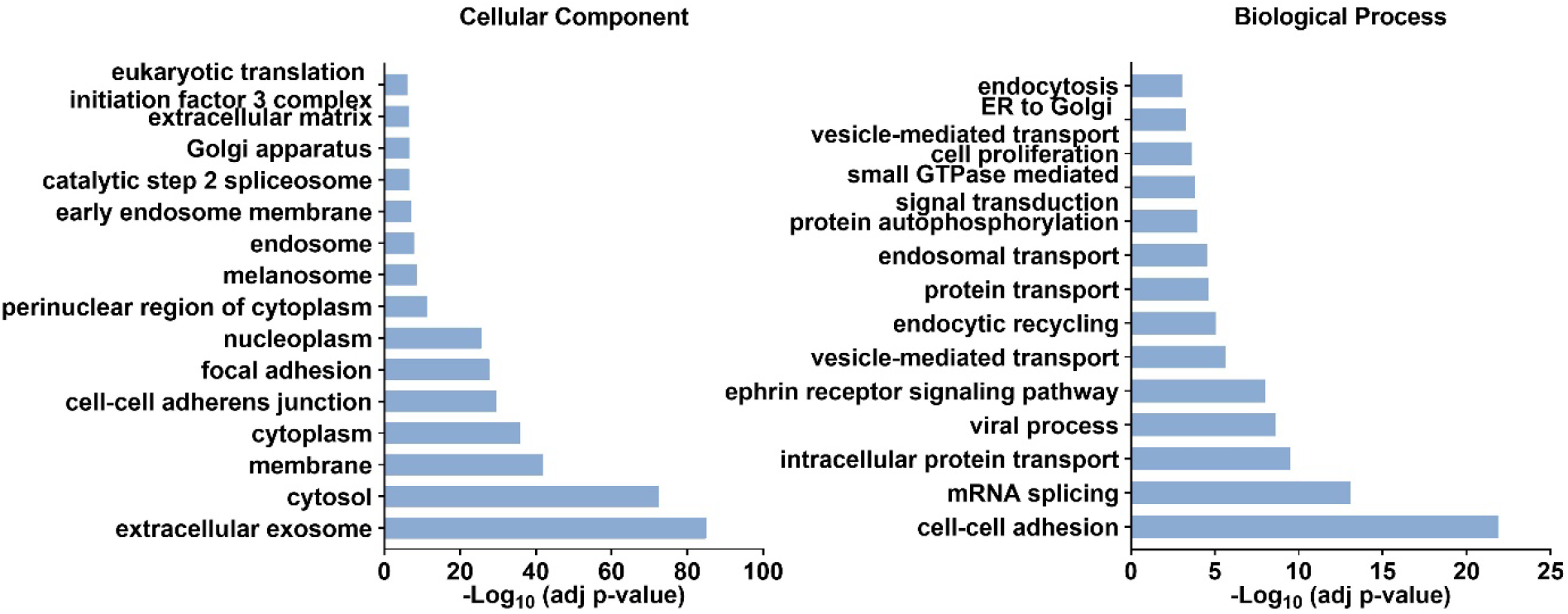
Gene ontology enrichment analysis of the “core” proteins commonly identified in EVs from all cell lines based on their cellular localisation and biological processes using the DAVID database and applying adjusted p-value < 0.001. For the cellular component top 15 significantly enriched terms are shown, full list of the terms can be found in the Supplementary Data.

We next investigated differences between the three cell lines using the 6 EV preparations as biological replicates. Unsupervised clustering and Principal Component Analysis (PCA) using the abundances of all detectable proteins clustered the cell lines separately, which was expected considering their inherent differences in molecular subtypes [50, 51]. The BT-474 EVs clustered more closely with the MCF7 EV preparations (PCA plot). It was also clear there was the most variability, in principal component 1, within the 6 EV preparations for BT-474, perhaps indicating a change in the cellular growth conditions (**Fig. 8A**). Supporting this, EV production dropped in the latter 3 timepoints with fewer particles (from 42-126⨯10^9^ down to 7-19⨯10^9^ particles/ml) and protein (from 147-365 µg down to 19-26 µg) than those collected prior to what we have termed ‘a catastrophic event’ between samples 10 and 11 (**Fig. 4A**).

**Figure 8.**
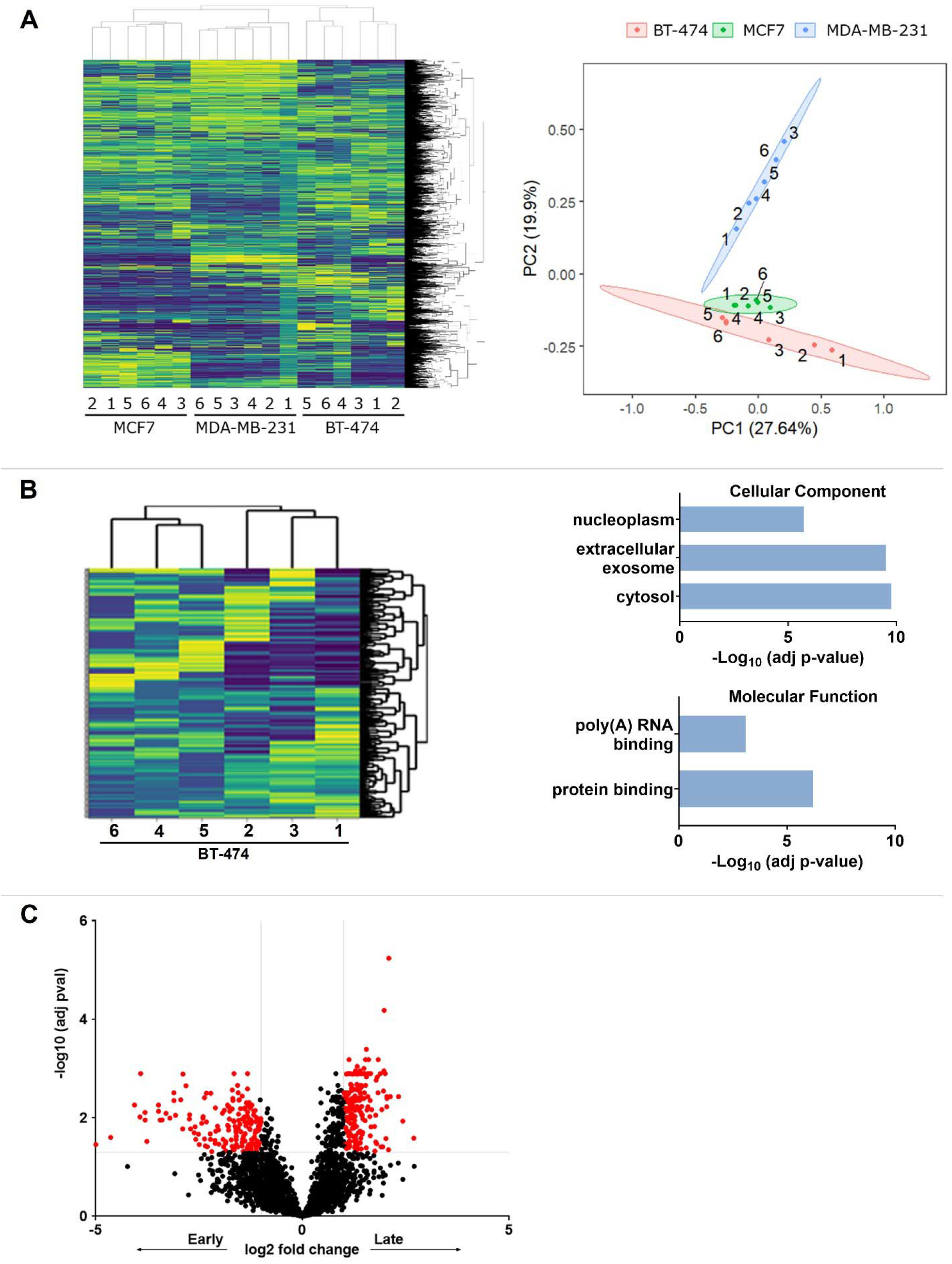
(A) Heatmap of all 2,875 detected proteins across all EV samples. Protein abundance was Z-score normalised and subjected to unsupervised clustering. Each row is a protein; columns represent separate experimental samples. Yellow, higher level of expression; blue, low level of expression. Principal component analysis (PCA) based on all proteins detected in EV samples. Plots show the first two principal components and their relative contribution to overall variance. (B) Identification of the most variable EV protein cargos in BT-474 EV samples. Heat map shows the sample clustering using the expression of the most variable proteins (SD cut off 1.0). The gene ontology categorisation analysis of most variable proteins is based on their cellular component and molecular function using DAVID software (adjusted p-value cut-off 0.001). (C) Volcano plot shows the comparison between BT-474 time points – Late and Early time points. The horizontal line corresponds to the adjusted p-value cut-off of 0.05. The two vertical lines show the positive and negative fold changes that equal 1. Dots in red represent proteins that are significantly abundant with a fold change greater than one.

Pairwise comparisons between the three cell lines identified relatively enriched proteins in their EVs. Gene ontology analysis of the differentially abundant EV proteins identified interesting biological processes and molecular functions reflective of the known characteristics of the parent cell line. For example, EVs derived from MDA-MB-231 cells, a triple negative line, were enriched for proteins involved in extracellular matrix organisation and disassembly through integrin and cadherin binding, cell adhesion, leukocyte migration and angiogenesis relative to the EV protein from MCF7 and BT-474. Conversely, EVs from luminal subtype MCF7 and BT-474 were enriched for translation processes, ribosomes, RNA binding and signalling relative to MDA-MB-231 EVs (Suppl. Fig. 6 and Suppl. Data).

Next, we asked the novel and important question of whether the 6 EV preparations taken during the bioreactor lifetime carry a consistent protein cargo and, if not, what external factors could lead to changes. To do this, we assessed the most variable proteins by abundance between the 6 replicates within each line by applying a relative Standard Deviation cut off of 1.0. Interestingly for MDA-MB-231 and MCF7, only 78 and 100 out of the 2,875 unique proteins respectively fit these criteria, representing no significantly enriched GO terms, highlighting that the EV protein cargo was relatively stable during the bioreactor lifetime. EV preparations from the BT-474 bioreactor, however, had 251 proteins fitting the criteria and were enriched for GO biological processes including gene expression, mRNA splicing and canonical glycolysis, likely driven by the aforementioned catastrophic event (**Fig. 8B**). Interestingly, the most variable proteins associated with the EVs from the BT474 bioreactor were more often localised to the cellular cytosol (37.1%) than to extracellular exosomes (33.1%), perhaps indicating cell death linked to the ‘catastrophic event’. We identified just 15 common ‘most variable’ proteins across all cell lines. To further investigate the stability of the EV protein cargo over the lifetime of the bioreactors we compared the 1,191 core proteins to the ‘most variable’ protein lists finding that 92.6% (1,103) did not overlap, highlighting the consistency in packaging of the core EV protein cargo throughout the extended lifespan of a bioreactor system (Suppl. Fig. 7).

In our final analyses, we investigated our hypotheses that the environmental conditions within the bioreactors can change the EV proteome. First, the day of the week for EV collection (Thursday Vs Monday) might alter EV cargo due to nutrient stress and second, that the EV protein cargo might change with the life of the bioreactor, differing between the three earliest and three latest EV preparations to reflect the changing growth conditions as the 3D cell structure is established and ages. To assess this, we performed pairwise comparisons of these subgroups within each cell line.

Based on our analysis, no proteins were differentially abundant due to either feeding cycle or time of bioreactor growth in MCF7 and MDA-MD-231 lines. Overall, these two bioreactors produced highly consistent EVs during their lifetimes. Feeding cycle did not affect protein cargo in the BT-474 derived EVs either. However, timepoints (Early and Late) affected the cargo of EV proteins from this cell line. Thus, 496 out of 2,875 unique proteins were differentially abundant between the Early and Late EV preparations (**Fig. 8C**). GO biological process analysis of these highlighted enrichment for cell-cell adhesion as well as mRNA splicing and processing in the early EV preparations. EV proteins from later timepoints were enriched for diverse biological process, such as intracellular protein transport, RNA splicing, antigen processing, endocytosis (Suppl. Data). This effect and the high number of overall variable proteins in the BT-474 cell line EV preparations (**Fig. 8**) is again likely due to the catastrophic event occurring between proteomic EV preparations 3 and 4 (NTA samples 10 and 11 Fig. 4A).

## Discussion

Many benefits are already well-reported in the literature for the use of bioreactor systems for large scale EV production [16, 21, 22, 25, 26, 33, 43, 52, 53]. For many researchers, the lower serum usage, removal of concentration steps, and incredible reduction of manual labour makes these systems highly cost and resource effective. Practically, the CELLine adherent bioreactor flasks have many advantages compared to producing EVs using conventional tissue culture flasks. For example, limiting the EV production to the 15 mL cell culture chamber removes the need for tangential flow and centrifugal concentrators prior to EV isolation which are known to lead to significant EV loss [54-57]. In addition, cells do not need to be passaged regularly as they naturally turn over during their continuous cultures which adds convenience for holidays and, of current relevance, hindered laboratory access. Furthermore, we have shown through imaging, that the cells grow as 3D structures, which is more physiologically relevant [58, 59], particularly for cancer, than growth of cells on tissue culture polystyrene as 2D monolayers.

The production of EVs that are free of contaminating EVs from serum supplementation is essential for many downstream applications. However, while commercially EV-depleted serum is available, it can be cost prohibitive or create challenges by the need to import restricted non-local animal-derived material. In addition, in-house depletion methods via ultracentrifugation and filtration are not capable of removing all of the exogenous EVs [60]. The CELLine bioreactors avoid these issues by allowing FBS-containing media to be used in the upper media chamber with contaminant bovine EVs excluded from entering the lower cell conditioned media by the 10 kDa filter membrane [20]. However, we recommend seeding the cells in serum-replete media to support adhesion to the growth surface prior to steady conversion of the media into serum depleted media to then commence EV collection. In addition, supplementation with CDM-HD serum replacement supported the high-density cell growth of all of the cell lines tested, allowing the continual collection of 24 to 37 EV harvests in this study, although whether this additive is essential is unclear. To date we have successfully cultured a number of immortalised cells and cancer cell lines using these conditions [52].

One of the disadvantages of this bioreactor system is that the cells cannot be visualised during growth. As part of this study, we aimed to develop a protocol for surrogate monitoring of bioreactor health. First, we considered non-invasively monitoring the cell chamber media volume and by counting and assessing the viability of the shed cells. In our analyses it was clear that although the former was able to identify bioreactors with a damaged membrane, the latter was too variable for interpretation. In fact, the three bioreactor replicates for the BT474 cell line each exhibited a unique cell shedding and viability pattern. The manufacturers of the CELLine-AD 1000 state that bioreactors have a maximum capacity of 4×10^8^ live cells and so the shed cell pattern may be highly dependent on the seeding density, as well as changing growth rates due to a range of environmental factors that cannot be well-controlled in a laboratory incubator. The indirect monitoring of the bioreactors through assessment of cell media volume and EV particle counts, did however prove to be a somewhat effective surrogate marker of bioreactor health. We therefore recommend that cell chamber media volume is used as a surrogate read-out of bioreactor health, particularly for assessing the integrity of the semi-permeable dialysis membrane, alongside EV particle number/protein analysis.

Deconstruction of the bioreactors following termination of the cultures allowed effective imaging of the growth surfaces by SEM and sectioning followed by H&E staining for imaging. To the best of our knowledge, this is the first example of analysing the 3D cell structure for cells grown in CELLine-AD bioreactors. The drastic increase in EV production relative to conventional flasks appears to be due largely to the thickness of cell masses and increased cell numbers, presumably by orders of magnitude as compared to conventional monolayer culture. The 3D structures also offer organisation of the cells into more tissue-like arrangements which is likely to alter their biology and the cargo and activities of the released EVs. Compared to conventional cultures, the bioreactors produced vast quantities of EVs, equivalent of up to 132x T175 flasks for the two lines directly assessed in this study. The effects of growth conditions on inducing EV release has been reported previously, for example for gastric cancer cells grown as spheroids [61], or MDA-MB-231 cells grown on collagen or matrigel coated fibre matrix [62] or 1D fibre-mimetic microtracks [14]. Interestingly our SEC-purified bioreactor EVs had more protein cargo per EV when compared to EVs from cells grown as monolayers. A reason for this discrepancy could be the increased cell turnover, leading to concentration of debris or the induction of free protein secretion due to growth conditions within the bioreactors, ultimately causing the formation of a protein corona coating EVs in the media. The 3D growth could also have modified the subtypes of EVs and mechanisms of release leading to a different and/or overall increased protein cargo. The importance of EV protein coronas is being increasingly reported, particularly in biofluids such as plasma that are protein rich. One recent study highlighted the association of extracellular matrix, complement system, immunoglobulins, coagulation factors, lipoproteins, nucleic acids, and thiollJreactive antioxidants coating cell line derived EVs spiked into plasma [63]. Components of the corona can therefore be characteristic of the surrounding matrix and it is proposed to be formed by protein aggregation and electrostatic interactions with EVs [64].

Our in-depth comparison of EV protein profiles using SWATH-MS from three breast cancer cell lines grown in bioreactor flasks identified common EV-associated proteins involved in cell adhesion and proliferation, key to the metastatic process, and other known breast cancer-associated biology. We defined a “core” of 1,191 EV proteins, present in every sample, which include known EV proteins such as integrins, annexins, and cytoskeletal proteins that mediate EV formation, facilitate binding to recipient cells, and direct fusion with recipient cell plasma membranes [65-69]. These proteins were significantly enriched for those associated with selected cellular components; ‘extracellular exosome’, ‘membrane’, ‘cytoplasm’ and ‘cell-cell adherens junction’. The enriched biological process terms for these core EV proteins included many cancer-relevant physiological and pathological processes, including cell-cell adhesion, mRNA splicing, and intracellular protein transport.

To further assess the concept of 3D growth in the bioreactor altering EV protein cargo [70], we performed a comparison to a published proteomics dataset of EVs, isolated using simple differential centrifugation, from MDA-MB-231 and MCF7 grown as monolayers [71]. Quantitative comparison was not feasible due to differences in proteomics methodology so a simple comparison of their 100 most abundant EV proteins in each cell line/condition with those identified herein was performed. Common proteins to both 2D and 3D grown EVs from both cell lines were enriched in GO molecular functions for cell adhesion molecular binding, cadherin binding and chaperones. Interestingly, the EVs from 2D cultures were enriched in RNA and carbohydrate derivative binding proteins, as well as histones, perhaps indicative of the presence of contaminating free proteins in the EV preparations, as well as proteins localised to focal adhesions, cell and anchoring junctions, structures that might be lacking in the 3D cultures to allow cell migration [72]. These differences suggest that the 3D high-density culture affects EV cargo, however, a direct comparison is needed to confirm these assumptions. It would also be interesting to assess whether the growth conditions affect the function of the EVs as well as their cargo, as recently reported for mesenchymal stem cells [19, 73]. Furthermore we note that although high culture cell viability is recommended for EV preparations in the MISEV2018 guidelines [45], the conditions in the bioreactor system may be more physiologically relevant due to the 3D structure and natural cell turnover which is abundant in tumours.

Unsupervised clustering of differentially expressed proteins between three cell lines revealed numerous subtype-specific protein clusters that reflect the known biology of breast cancer molecular subtypes. EV proteins from a clinically more aggressive triple negative subtype, MDA-MB-231, were associated with angiogenesis, cell motility and migration and were more different from other two hormone receptor positive cell lines – MCF7 and BT-474. Within each cell line, bioreactor replicates clustered together, demonstrating the technical and biological reproducibility of EV isolation, bioreactor growth, and robust SWATH-MS quantification. The exception was for BT-474, where EV protein cargo segregated the preparations from earlier from later stages of bioreactor culture. Comparison of most variable proteins from all three cell lines with the identified 1,191 ‘core’ proteins importantly highlighted the stability of the majority of these proteins over the lifetime of bioreactors. Moreover, the EV proteomes from MDA-MB-231 and MCF7 bioreactors were not on the whole affected by the feeding cycle nor by the timepoint of EV collection, demonstrating an overall high consistency of EV protien content during the bioreactor lifetimes. One of the reasons for the minimal variability in the bioreactor EV preparations may be due to allowing 4 weeks for the system to establish during the conversion to serum-free media, stabilising the 3D growth structures and reaching a steady state, prior to EV sampling.

Although not quantitated in our SWATH-MS data, the tetraspanin profiles of pooled bioreactor EVs of the five breast-cancer cell lines differed by immunoblot. With the recent report using HeLa cells of CD63 being more frequently co-localised with exosomes and conversely CD9 and CD81 more likely to be associated with surface membrane ectosomes [74], this highlights potential differences in EV biogenesis mechanisms between molecular subtypes of breast cancer. Limited EV-associated CD63 and abundant CD81 points towards cell budding EV formation in the SKBR3 and BT-474 bioreactors that we hypothesise underwent ‘catastrophic events’ based on EV particle number, therefore may be indicative of initiation of various regulated cell death mechanisms [75]. SEM of the terminated bioreactors also found fewer cells on the growth surface for these two cell lines. The catastrophic event was not investigated further in the SKBR3 cells but we note that curiously the EV preparations from this line consistently co-isolated with white, lipid rich material, a feature not before reported in the literature. Cell growth environment has been reported to affect cellular function and morphology and it is known that breast cancer cells can contribute to normal mammary gland development by retaining the ability to differentiate and secrete milk proteins [76]. In particular, HER2+ cell lines, like SKBR3, rely on a Warburg-like metabolism for survival and invasive behaviour, dependent on fatty acid synthesis with increased storage of fat droplets. Overexpression of HER2 has pro-lipogenic effects and these cells are shown to store more triacylglycerides and saturated fatty acids [77, 78]. Therefore, we speculate that the bioreactor system may have induced these processes towards differentiation of the ductal breast cancer cells to produce milk products that were co-isolated with our EV preparations.

A reduction in BT-474 EV number occurred between proteomics preparations 3 and 4 (NTA samples 10 and 11) where we detected concomitant changes to the protein cargo by SWATH. GO of functions changed from cell adhesion and mRNA splicing and processing, to protein transport and vesicle-mediated transport, ephrin receptor signalling pathway, antigen processing, endocytosis in the later EV cargos. The pre-catastrophe EV cargo proteins reflect normal processes occurring in the BT-474 cells, for example cell-cell adhesion that would be evident in a minimally invasive cell-line [79]. The exact cause of the catastrophe is not evident from any of our data. We note that all cell lines were tested for mycoplasma prior to seeding and as part of our maintenance schedule particularly turbid cell media was streaked onto agar plates to assess for bacterial contamination. However, it is possible that the use of antibiotics and antimycotics plus the long-term growth could have led to a low-level infection in the system. This ‘catastrophic event’ was not repeated with the replicate two BT-474 bioreactors, but also happened to SKBR3, another HER2+ line, highlighting the importance of carefully monitoring bioreactor health using EV number to avoid major differences in the quality and cargo of the EV preparations.

## Conclusions

In this study, we have demonstrated that the CELLine AD 1000 bioreactor system allowed the consistent and simultaneous production of EVs from several breast cancer cell lines of different subtypes. Using the reported inoculation, monitoring, EV purification and characterisation protocols, researchers in many fields will be able to produce vast numbers of EVs with much less resource commitment compared to using conventional tissue culture flasks. In addition, the reported methods of imaging of the bioreactor growth surfaces may be useful in other applications to assess the growth characteristics of the cultured cells. Lastly, our finding that the EV yields, and protein cargo profiles are indicative of the cell of origin, and can indicate cellular health, gives confidence in the consistency of bioreactor samples while providing protocols for quality monitoring of EV preparations.

## Acknowledgements

The authors would like to thank the Breast Cancer Foundation New Zealand for supporting this work through a Technology and Innovation Project Grant. We would also like to thank the Hub for Extracellular Vesicle Investigations (HEVI) for their continued support and the University of Auckland’s Mass Spectrometry Hub (MaSH) for their assistance in the proteomics analysis.

---

**Supplementary Figure 1.**
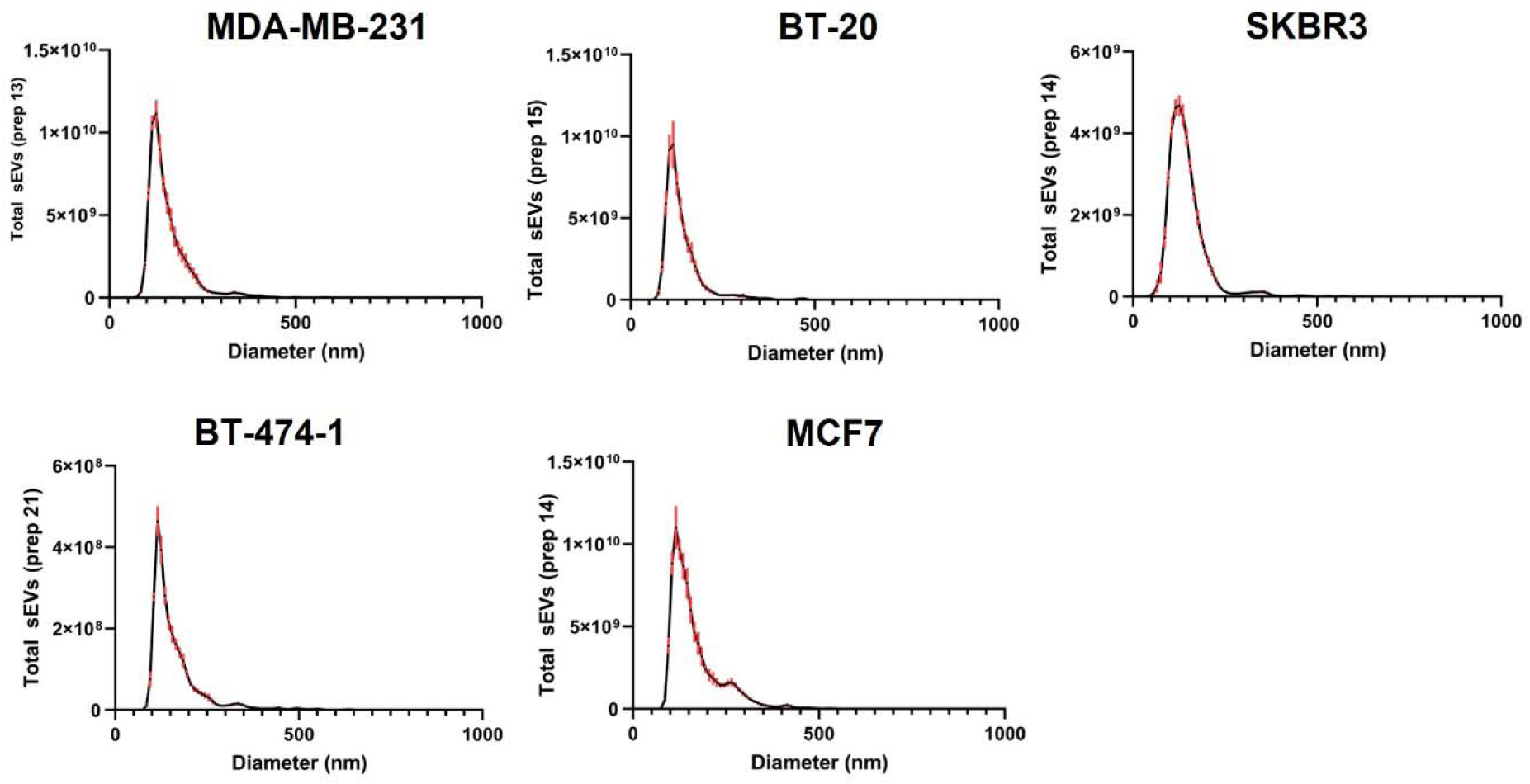
Particle size distribution profiles as determined by NTA. Data shown from representative EV preparations isolated from all bioreactors.

**Supplementary Figure 2.**
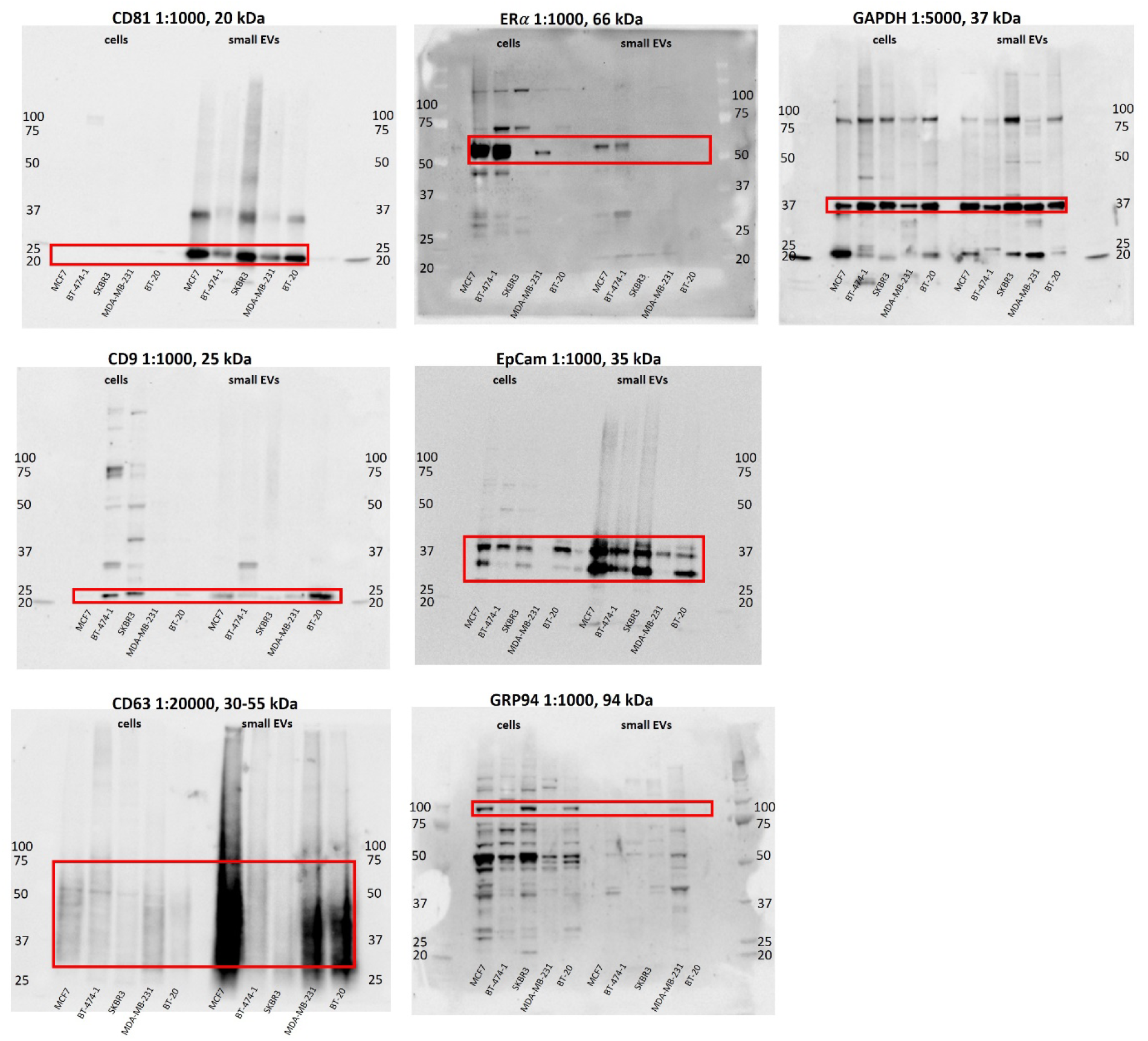
Uncropped blot images. Abundance of CD81, CD9, CD63, ERα, EpCAM, GRP94 and GAPDH in MCF7, BT-474-1, SKBR3, MDA-MB-231, BT-20 whole cell lysates and small EVs.

**Supplementary Figure 3.**
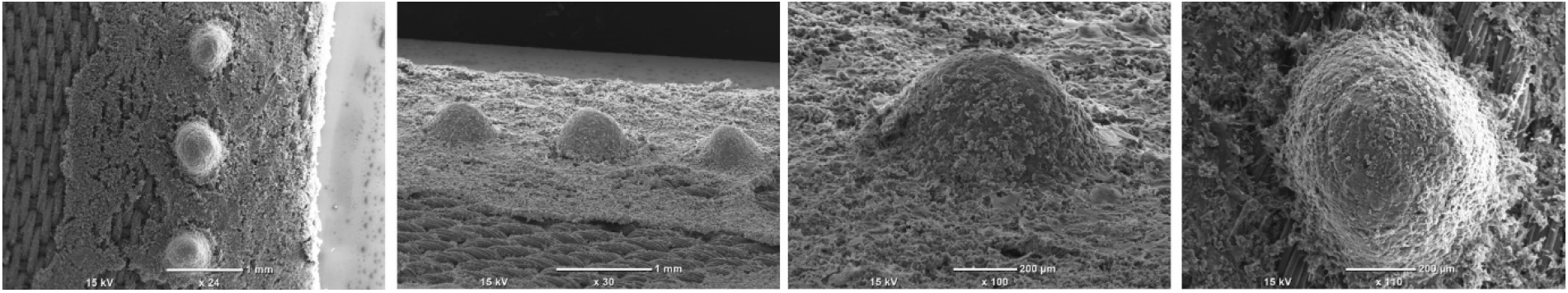
Spheroid like cell masses on the topside of the growth surface of the BT-20 bioreactor. These structures were not seen in any of the other samples imaged. Scale bars 1 mm and 200 µm.

**Supplementary Figure 4.**
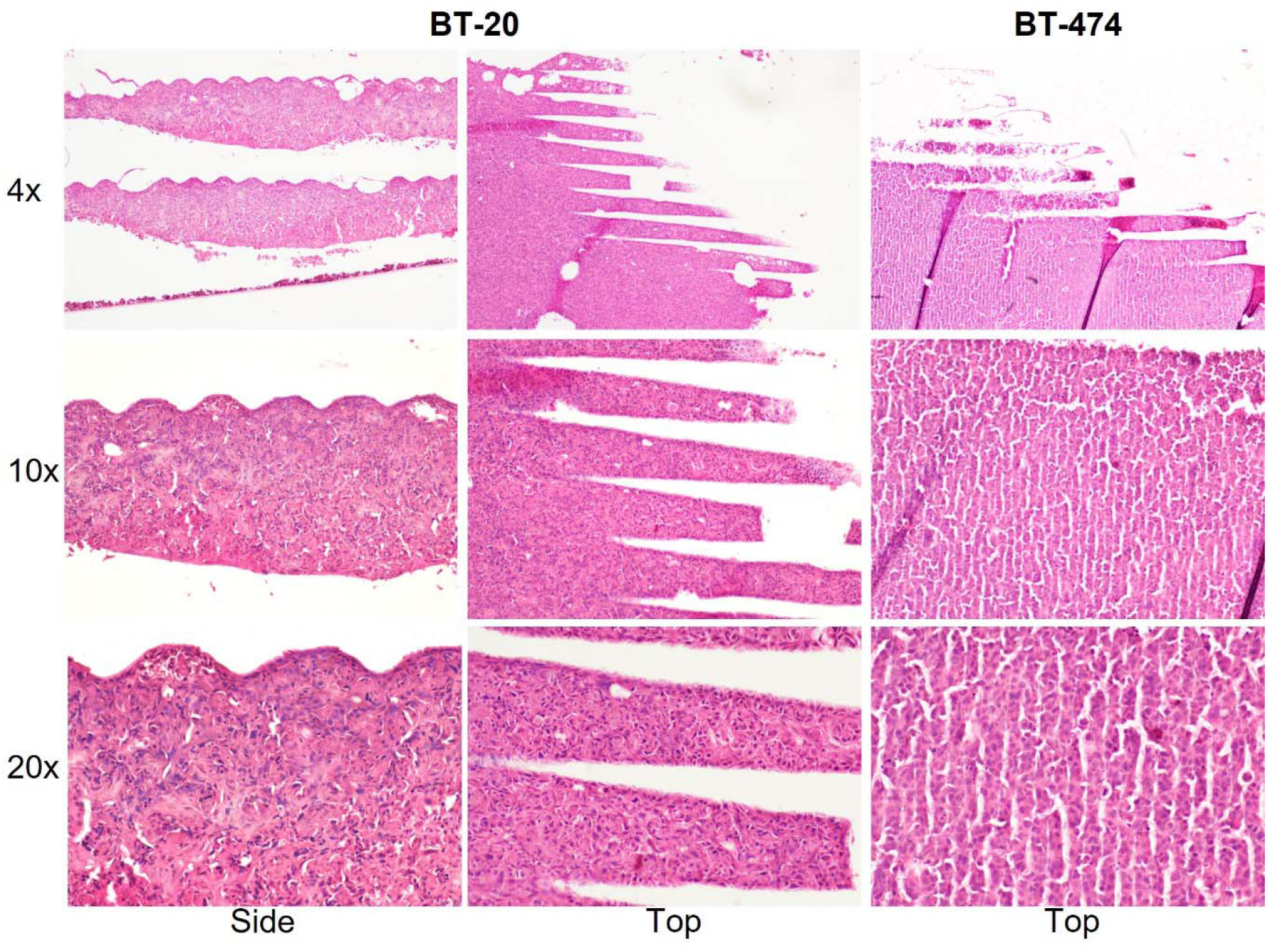
Representative images showing H&E staining of sections of BT-20 (left) and BT-474 (right) bioreactor growth surface. The side and top view is shown for BT-20 bioreactor, the top view is shown for the BT-474 bioreactor. The magnification of the images is shown on the left hand side.

**Supplementary Figure 5.**
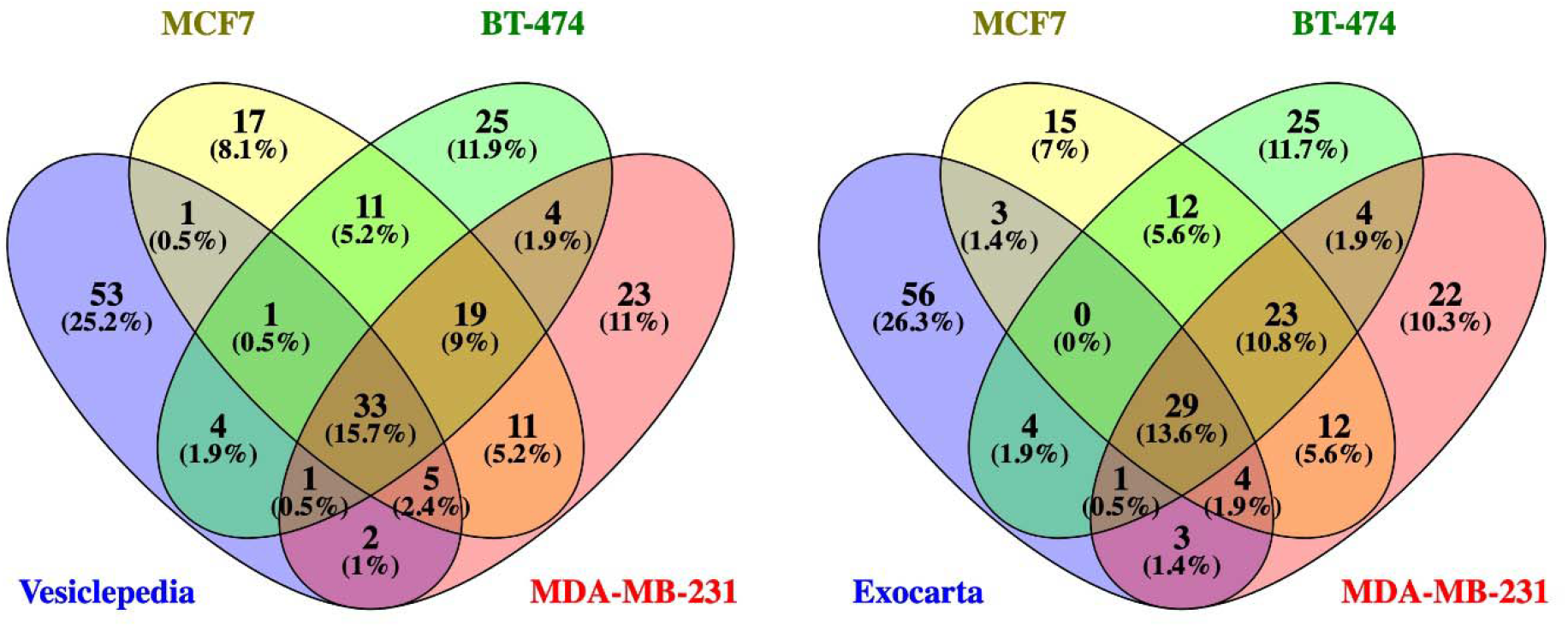
Venn diagrams showing overlap of the top 100 most abundant proteins identified within MCF7, BT-474, and MDA-MB-231 small EVs and the top 100 proteins that are often identified in EVs in the (left) Vesiclepedia and (right) Exocarta databases.

**Supplementary Figure 6.**
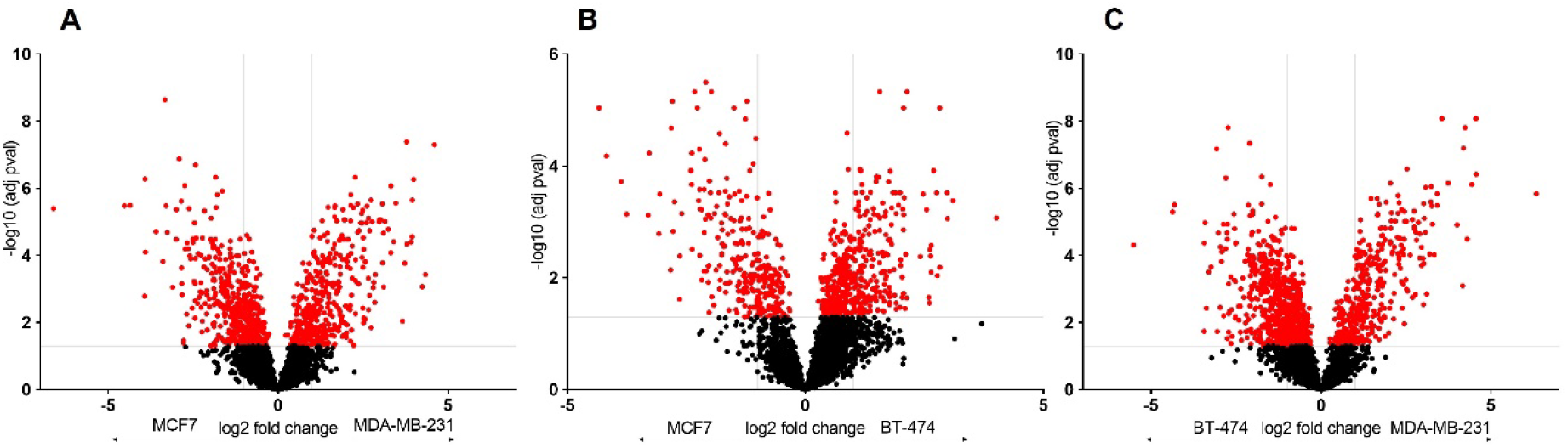
Volcano plots showing a pairwise comparison between cell lines: (A) MCF7 and MDA-MB-231, (B) MCF7 and BT-474; (C) BT-474 and MDA-MB-231. The horizontal line corresponds to the adjusted p-value cut-off of 0.05. The two vertical lines show the positive and negative fold changes that equal 1. Dots in red represent proteins that are significantly abundant based on the p-value.

**Supplementary Figure 7.**
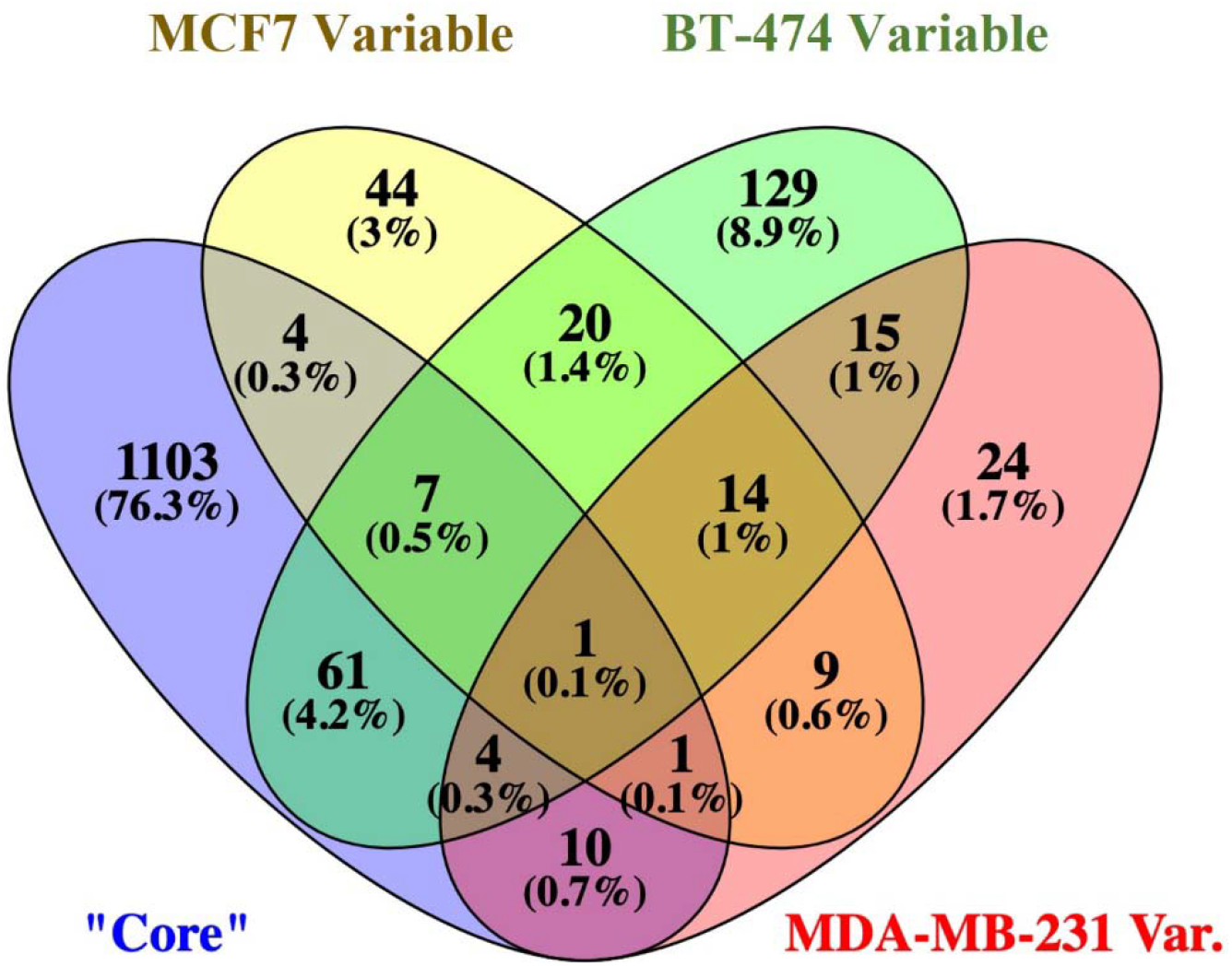
Comparison of the most variable proteins from EVs isolated from MCF7, BT-474 and MDA-MB-231 bioreactors with the “core” proteins.

## References

1. Margolis, L. and Y. Sadovsky, The biology of extracellular vesicles: The known unknowns. PLoS Biol, 2019. 17(7): p. e3000363.

2. Joshi, B.S., et al., Endocytosis of Extracellular Vesicles and Release of Their Cargo from Endosomes. ACS Nano, 2020. 14(4): p. 4444–4455.

3. Raposo, G. and W. Stoorvogel, Extracellular vesicles: exosomes, microvesicles, and friends. J Cell Biol, 2013. 200(4): p. 373–83.

4. Gowen, A., et al., Mesenchymal Stem Cell-Derived Extracellular Vesicles: Challenges in Clinical Applications. Front Cell Dev Biol, 2020. 8: p. 149.

5. Rani, S., et al., Mesenchymal Stem Cell-derived Extracellular Vesicles: Toward Cell-free Therapeutic Applications. Mol Ther, 2015. 23(5): p. 812–823.

6. Park, K.S., et al., Enhancement of therapeutic potential of mesenchymal stem cell-derived extracellular vesicles. Stem Cell Res Ther, 2019. 10(1): p. 288.

7. Zhang, Z., J.A. Dombroski, and M.R. King, Engineering of Exosomes to Target Cancer Metastasis. Cell Mol Bioeng, 2020. 13(1): p. 1–16.

8. Murphy, D.E., et al., Extracellular vesicle-based therapeutics: natural versus engineered targeting and trafficking. Exp Mol Med, 2019. 51(3): p. 1–12.

9. Piffoux, M., et al., Modification of Extracellular Vesicles by Fusion with Liposomes for the Design of Personalized Biogenic Drug Delivery Systems. ACS Nano, 2018. 12(7): p. 6830–6842.

10. Zhou, B., et al., Application of exosomes as liquid biopsy in clinical diagnosis. Signal Transduct Target Ther, 2020. 5(1): p. 144.

11. Yu, W., et al., Exosome-based liquid biopsies in cancer: opportunities and challenges. Ann Oncol, 2021. 32(4): p. 466–477.

12. Armstrong, D. and D.E. Wildman, Extracellular Vesicles and the Promise of Continuous Liquid Biopsies. J Pathol Transl Med, 2018. 52(1): p. 1–8.

13. Chen, Y.S., et al., Exosomes in clinical trial and their production in compliance with good manufacturing practice. Ci Ji Yi Xue Za Zhi, 2020. 32(2): p. 113–120.

14. Hisey, C.L., et al., Micropatterned growth surface topography affects extracellular vesicle production. Colloids Surf B Biointerfaces, 2021. 203: p. 111772.

15. Patel, D.B., et al., Towards rationally designed biomanufacturing of therapeutic extracellular vesicles: impact of the bioproduction microenvironment. Biotechnol Adv, 2018. 36(8): p. 2051–2059.

16. Yan, L. and X. Wu, Exosomes produced from 3D cultures of umbilical cord mesenchymal stem cells in a hollow-fiber bioreactor show improved osteochondral regeneration activity. Cell Biol Toxicol, 2020. 36(2): p. 165–178.

17. Ouyang, B., et al., Extracellular Vesicles From Human Urine-Derived Stem Cells Ameliorate Erectile Dysfunction in a Diabetic Rat Model by Delivering Proangiogenic MicroRNA. Sex Med, 2019. 7(2): p. 241–250.

18. Maji, S., et al., In vitro toxicology studies of extracellular vesicles. J Appl Toxicol, 2017. 37(3): p. 310–318.

19. Cao, J., et al., Three-dimensional culture of MSCs produces exosomes with improved yield and enhanced therapeutic efficacy for cisplatin-induced acute kidney injury. Stem Cell Res Ther, 2020. 11(1): p. 206.

20. Zoe, T., et al., Scale-Up Production Of Extracellular Vesicles From Mesenchymal Stromal Cells Isolated From Pre-Implantation Equine Embryos. Scientific Reports, 2022.

21. Watson, D.C., et al., Efficient production and enhanced tumor delivery of engineered extracellular vesicles. Biomaterials, 2016. 105: p. 195–205.

22. Watson, D.C., et al., Scalable, cGMP-compatible purification of extracellular vesicles carrying bioactive human heterodimeric IL-15/lactadherin complexes. J Extracell Vesicles, 2018. 7(1): p. 1442088.

23. Yoo, K.W., et al., Large-Scale Preparation of Extracellular Vesicles Enriched with Specific microRNA. Tissue Eng Part C Methods, 2018. 24(11): p. 637–644.

24. Patel, D.B., et al., Enhanced extracellular vesicle production and ethanol-mediated vascularization bioactivity via a 3D-printed scaffold-perfusion bioreactor system. Acta Biomater, 2019. 95: p. 236–244.

25. Mitchell, J.P., et al., Increased exosome production from tumour cell cultures using the Integra CELLine Culture System. J Immunol Methods, 2008. 335(1-2): p. 98–105.

26. Guerreiro, E.M., et al., Efficient extracellular vesicle isolation by combining cell media modifications, ultrafiltration, and size-exclusion chromatography. PLoS One, 2018. 13(9): p. e0204276.

27. Chen, M., et al., Distinct shed microvesicle and exosome microRNA signatures reveal diagnostic markers for colorectal cancer. PLoS One, 2019. 14(1): p. e0210003.

28. Xu, R., et al., Surfaceome of Exosomes Secreted from the Colorectal Cancer Cell Line SW480: Peripheral and Integral Membrane Proteins Analyzed by Proteolysis and TX114. Proteomics, 2019. 19(8): p. e1700453.

29. Ji, H., et al., Deep sequencing of RNA from three different extracellular vesicle (EV) subtypes released from the human LIM1863 colon cancer cell line uncovers distinct miRNA-enrichment signatures. PLoS One, 2014. 9(10): p. e110314.

30. Haug, B.H. OH;, Utnes, P; Roth, SA; Lokke, C; Flaegstad, T; Einvik, C, Exosome-like EVs from MYCN-amplified neuroblastoma cells contain oncogenic miRNAs. Anticancer Research, 2015. 35: p. 2521–2530.

31. Griffiths, S.G., et al., Differential Proteome Analysis of Extracellular Vesicles from Breast Cancer Cell Lines by Chaperone Affinity Enrichment. Proteomes, 2017. 5(4).

32. Jeppesen, D.K., et al., Quantitative proteomics of fractionated membrane and lumen exosome proteins from isogenic metastatic and nonmetastatic bladder cancer cells reveal differential expression of EMT factors. Proteomics, 2014. 14(6): p. 699–712.

33. Hisey, C.L., et al., Towards establishing extracellular vesicle-associated RNAs as biomarkers for HER2+ breast cancer. F1000Res, 2020. 9: p. 1362.

34. Faruqu, F.N., L. Xu, and K.T. Al-Jamal, Preparation of Exosomes for siRNA Delivery to Cancer Cells. J Vis Exp, 2018(142).

35. Palviainen, M., et al., Metabolic signature of extracellular vesicles depends on the cell culture conditions. J Extracell Vesicles, 2019. 8(1): p. 1596669.

36. Green, T.M., et al., Breast Cancer-Derived Extracellular Vesicles: Characterization and Contribution to the Metastatic Phenotype. Biomed Res Int, 2015. 2015: p. 634865.

37. Sadovvska, L., J. Eglitis, and A. Line, Extracellular Vesicles as Biomarkers and Therapeutic Targets in Breast Cancer. Anticancer Research, 2015. 35: p. 6379–90.

38. Wang, H.X. and O. Gires, Tumor-derived extracellular vesicles in breast cancer: From bench to bedside. Cancer Lett, 2019. 460: p. 54–64.

39. Keklikoglou, I., et al., Chemotherapy elicits pro-metastatic extracellular vesicles in breast cancer models. Nat Cell Biol, 2019. 21(2): p. 190–202.

40. Peng, J., et al., Roles of Extracellular Vesicles in Metastatic Breast Cancer. Breast Cancer (Auckl), 2018. 12: p. 1178223418767666.

41. Dai, X., et al., Breast Cancer Cell Line Classification and Its Relevance with Breast Tumor Subtyping. J Cancer, 2017. 8(16): p. 3131–3141.

42. CELLine User Manual. Available from: https://www.sigmaaldrich.com/deepweb/assets/sigmaaldrich/product/documents/374/057/celline-user-manual.pdf.

43. Mittermaier, J.Z.-G., M.O., Long-Term High Level Protein Expression in Adherent Protein-free Growing BHK Cells Using Integra CELLine Adhere Bioreactor Flasks. 2004: Genetic Engineering News. p. 42.

44. Jackson, L.R.T. L.,; Fox, J.G.; Lipman, N.S., Evaluation of hollow fiber bioreactors as an alternative to murine ascites production for small scale monoclonal antibody production. Journal of Immunological Methods, 1996. 189: p. 217–231.

45. Thery, C., et al., Minimal information for studies of extracellular vesicles 2018 (MISEV2018): a position statement of the International Society for Extracellular Vesicles and update of the MISEV2014 guidelines. J Extracell Vesicles, 2018. 7(1): p. 1535750.

46. Simpson, R.J., H. Kalra, and S. Mathivanan, ExoCarta as a resource for exosomal research. J Extracell Vesicles, 2012. 1.

47. Kalra, H., et al., Vesiclepedia: a compendium for extracellular vesicles with continuous community annotation. PLoS Biol, 2012. 10(12): p. e1001450.

48. Kowal, J., et al., Proteomic comparison defines novel markers to characterize heterogeneous populations of extracellular vesicle subtypes. Proc Natl Acad Sci U S A, 2016. 113(8): p. E968–77.

49. Huang da, W., B.T. Sherman, and R.A. Lempicki, Systematic and integrative analysis of large gene lists using DAVID bioinformatics resources. Nat Protoc, 2009. 4(1): p. 44–57.

50. Dai, X.L. T.,; Bai, Z.; Yang, Y., Liu, X., Zhan, J., Shi, B., Breast cancer intrinsic subtype classification -clinical use and future trends. Am J of Cancer Research, 2015. 5(10): p. 2929–2943.

51. Yersal, O. and S. Barutca, Biological subtypes of breast cancer: Prognostic and therapeutic implications. World J Clin Oncol, 2014. 5(3): p. 412–24.

52. Artuyants, A.C. V.; Reshef, V.; Blenkiron, C.; Chamley, L.W.; Leung, E.; Hisey, C.L., Production of Extracellular Vesicles Using a CELLine Adherent Bioreactor Flask. Methods in Molecular Biology, 2021.

53. Yan, I.S., N; Borrelli, DA; Patel, T, Use of a Hollow Fiber Bioreactor to Collect EVs from Cells in Culture. Extracellular RNA: Methods in Molecular Biology, ed. J. Walker. Vol. 1740. 2018: Springer Protocols.

54. Li, P., et al., Progress in Exosome Isolation Techniques. Theranostics, 2017. 7(3): p. 789–804.

55. Ramirez, M.I., et al., Technical challenges of working with extracellular vesicles. Nanoscale, 2018. 10(3): p. 881–906.

56. van de Wakker, S.I., et al., Influence of short term storage conditions, concentration methodsand excipients on extracellular vesicle recovery and function. European Journal of Pharmaceutics and Biopharmaceutics, 2022. 170: p. 59–69.

57. Vergauwen, G., et al., Confounding factors of ultrafiltration and protein analysis in extracellular vesicle research. Scientific Reports, 2017. 7(1): p. 2704.

58. Tu, J., et al., Cancer spheroids derived exosomes reveal more molecular features relevant to progressed cancer. Biochemistry and Biophysics Reports, 2021. 26: p. 101026.

59. Thippabhotla, S., C. Zhong, and M. He, 3D cell culture stimulates the secretion of in vivo like extracellular vesicles. Scientific Reports, 2019. 9(1): p. 13012.

60. Lehrich, B.M., Y. Liang, and M.S. Fiandaca, Foetal bovine serum influence on in vitro extracellular vesicle analyses. J Extracell Vesicles, 2021. 10(3): p. e12061.

61. Rocha, S., et al., 3D Cellular Architecture Affects MicroRNA and Protein Cargo of Extracellular Vesicles. Adv Sci (Weinh), 2019. 6(4): p. 1800948.

62. Franchi, M., et al., Extracellular Matrix-Mediated Breast Cancer Cells Morphological Alterations, Invasiveness, and Microvesicles/Exosomes Release. Cells, 2020. 9(9).

63. Toth, E.A., et al., Formation of a protein corona on the surface of extracellular vesicles in blood plasma. J Extracell Vesicles, 2021. 10(11): p. e12140.

64. Cedervall, T.L. I.,; Lindman, S.; Berggard, T.; Thulin, E.; Nilsson, H.; Dawson, K.A.; Linse, S., Understanding the nanoparticle-protein corona using methods to quantify exchange rates and affinities of proteins for nanoparticles. PNAS, 2007. 104(7): p. 2050–2055.

65. Altei, W.F., et al., Inhibition of αvβ3 integrin impairs adhesion and uptake of tumor-derived small extracellular vesicles. Cell Communication and Signaling, 2020. 18(1): p. 158.

66. Fuentes, P., et al., ITGB3-mediated uptake of small extracellular vesicles facilitates intercellular communication in breast cancer cells. Nature Communications, 2020. 11(1): p. 4261.

67. Koumangoye, R.B., et al., Detachment of Breast Tumor Cells Induces Rapid Secretion of Exosomes Which Subsequently Mediate Cellular Adhesion and Spreading. PLOS ONE, 2011. 6(9): p. e24234.

68. Tontanahal, A., I. Arvidsson, and D. Karpman, Annexin Induces Cellular Uptake of Extracellular Vesicles and Delays Disease in Escherichia coli O157:H7 Infection. Microorganisms, 2021. 9(6).

69. Blaser, M.C. and E. Aikawa, Roles and Regulation of Extracellular Vesicles in Cardiovascular Mineral Metabolism. Frontiers in Cardiovascular Medicine, 2018. 5.

70. Millan, C., et al., Extracellular Vesicles from 3D Engineered Microtissues Harbor Disease-Related Cargo Absent in EVs from 2D Cultures. Advanced Healthcare Materials, 2022. 11(5): p. 2002067.

71. Rontogianni, S., et al., Proteomic profiling of extracellular vesicles allows for human breast cancer subtyping. Commun Biol, 2019. 2: p. 325.

72. Harunaga, J.S. and K.M. Yamada, Cell-matrix adhesions in 3D. Matrix Biol, 2011. 30(7-8): p. 363–8.

73. Kusuma, G.D., et al., Effect of 2D and 3D Culture Microenvironments on Mesenchymal Stem Cell-Derived Extracellular Vesicles Potencies. Frontiers in Cell and Developmental Biology, 2022. 10.

74. Mathieu, M., et al., Specificities of exosome versus small ectosome secretion revealed by live intracellular tracking of CD63 and CD9. Nature Communications, 2021. 12(1): p. 4389.

75. Baxter, A.A., et al., Analysis of extracellular vesicles generated from monocytes under conditions of lytic cell death. Sci Rep, 2019. 9(1): p. 7538.

76. Bussard, K.M. and G.H. Smith, Human breast cancer cells are redirected to mammary epithelial cells upon interaction with the regenerating mammary gland microenvironment in-vivo. PLoS One, 2012. 7(11): p. e49221.

77. Baumann, J., C. Sevinsky, and D.S. Conklin, Lipid biology of breast cancer. Biochim Biophys Acta, 2013. 1831(10): p. 1509–17.

78. Kinlaw, W.B., et al., Fatty Acids and Breast Cancer: Make Them on Site or Have Them Delivered. J Cell Physiol, 2016. 231(10): p. 2128–41.

79. Fujita, M., et al., Metabolic characterization of aggressive breast cancer cells exhibiting invasive phenotype: impact of non-cytotoxic doses of 2-DG on diminishing invasiveness. BMC Cancer, 2020. 20(1): p. 929.

